# Frontotemporal cortex flexibly adapts latent structural representations

**DOI:** 10.64898/2026.06.25.734414

**Authors:** Georgios Tertikas, Nadescha Trudel, Miriam Klein-Flügge, Tobias U. Hauser

## Abstract

Humans excel at navigating complex environments by forming abstract structural representations that can be flexibly updated when environments change. Here, we examine how the brain dynamically reconfigures these internal models in response to covert changes in latent hierarchies. Using a novel inference task and fMRI repetition suppression, we find that stable relational knowledge is encoded in medial orbitofrontal cortex (mOFC), while structural changes trigger transient representations across hippocampal and prefrontal regions. Newly inferred associations are first encoded in anterior medial frontal cortex (amFC) and migrate ventrally to mOFC when settling. In contrast, outdated associations transiently engage frontopolar cortex and hippocampus, with the hippocampus, but not frontal areas, retaining a residual memory trace. Notably, the strength of early novel signals in amFC and hippocampus tracks individual differences in behavioural adaptation. Together, these results characterise mechanisms supporting adaptive structural reconfiguration in the human brain, with implications for cognitive inflexibility in psychiatric disorders.

**SIGNIFICANCE STATEMENT:** The brain relies on internal models of hidden environmental structure, yet how these models are revised when the world changes remains unclear. We developed a novel task and neuroimaging approach that allowed us to trace the emergence, maintenance, and dissolution of latent structural representations in humans. Stable representations were encoded in medial orbitofrontal cortex, whereas newly formed and outdated associations followed distinct temporal trajectories across frontal and hippocampal regions. Outdated representations transiently recruited frontopolar cortex but persisted in hippocampus after becoming behaviorally irrelevant. Neural signatures of updating predicted individual differences in adaptation, revealing candidate mechanisms through which cognitive flexibility, and its impairment in psychiatric disorders, may arise.

## INTRODUCTION

The world’s inner workings can often only be inferred. Whether you reach a destination on time depends not only on your explicit choice of route, but also on the inferred state of traffic. Predicting this state requires latent knowledge of the network,for instance, knowing what time is rush hour and which roads will be busy at that time. Such structural representations,mental models that organize and relate information,help to represent complex associations effectively (O’Reilly et al., 2022). Originally established for spatial navigation (Tolman, 1948; Maguire et al., 2000; Epstein et al., 2017), these latent structural representations are also used for abstract cognitive and social structures, enabling us to generalize knowledge and navigate abstract domains (Constantinescu et al., 2016; Park et al., 2020; Schwartenbeck et al., 2023; Cone & Clopath, 2024; Wittmann et al., 2025).

The brain relies on a clearly delineated fronto-hippocampal network to represent such latent structural associations. The hippocampus is critically involved in forming abstract representations and associations, mostly through encoding and retrieving spatial and episodic information (McKenzie et al., 2014; Barron et al., 2020; Garvert et al., 2023). These structural representations are subsequently represented in orbital and medial frontal cortex (OFC/mFC), where they are used to navigate in abstract spaces (Wilson et al., 2014; Schuck et al., 2016; Schuck et al., 2018; Klein-Flügge et al., 2019; Mızrak et al., 2021).

Whilst said network is well evidenced for building and representing latent structures (Bradfield et al., 2018; Bein & Niv, 2025), little is known about the mechanisms that allow established latent associations to be flexibly adjusted. When discovering that a primary bridge on a familiar route has been permanently closed, the latent structure of the road network—and the resulting behavioral policy—must be revised. In dynamic environments, flexible adaptation requires multiple processes: from detecting outdated associations to learning novel contingencies, whilst maintaining stable associations that have remained unchanged.

In this study, we investigated how changes to latent structural representations are flexibly and dynamically adapted in fronto-hippocampal networks. We combined a novel task design with analyses inspired byrepetition suppression (RS) analyses (Grill-Spector et al., 2006; Summerfield et al., 2008; Klein-Flügge et al., 2013; Barron et al., 2016; Boorman et al., 2016; Rustichini et al., 2017), leveraging the principle that neuronal responses are reduced when a stimulus is predicted or associated with prior experience. By measuring reduced neuronal signals between structurally related stimuli, we were able to assess the deprecation, re-formation, and maintenance of latent structural association. We find that these processes take place across distinct subregions of a fronto-hippocampal network, and that the neural code of these adaptations reflects the cognitive flexibility underlying participants’ ability to update hidden structural representations. Our findings provide insights relevant in the context of impaired cognitive flexibility in psychiatry patients (Robbins, 2007; Dajani & Uddin, 2015; McTeague et al., 2016; Shahar et al., 2021).

## RESULTS

### Hidden changes allow probing flexible adaptation of structural representations

To probe how humans learn and flexibly update latent structural representations, we trained participants (N=57; 51 in the imaging analysis, age: 25.6 ± 5.6 years, range 18-40 years, 23 male; see methods for training procedure and exclusion criteria) on a novel latent, hierarchical structure task (Figure 1).

**Figure 1.**
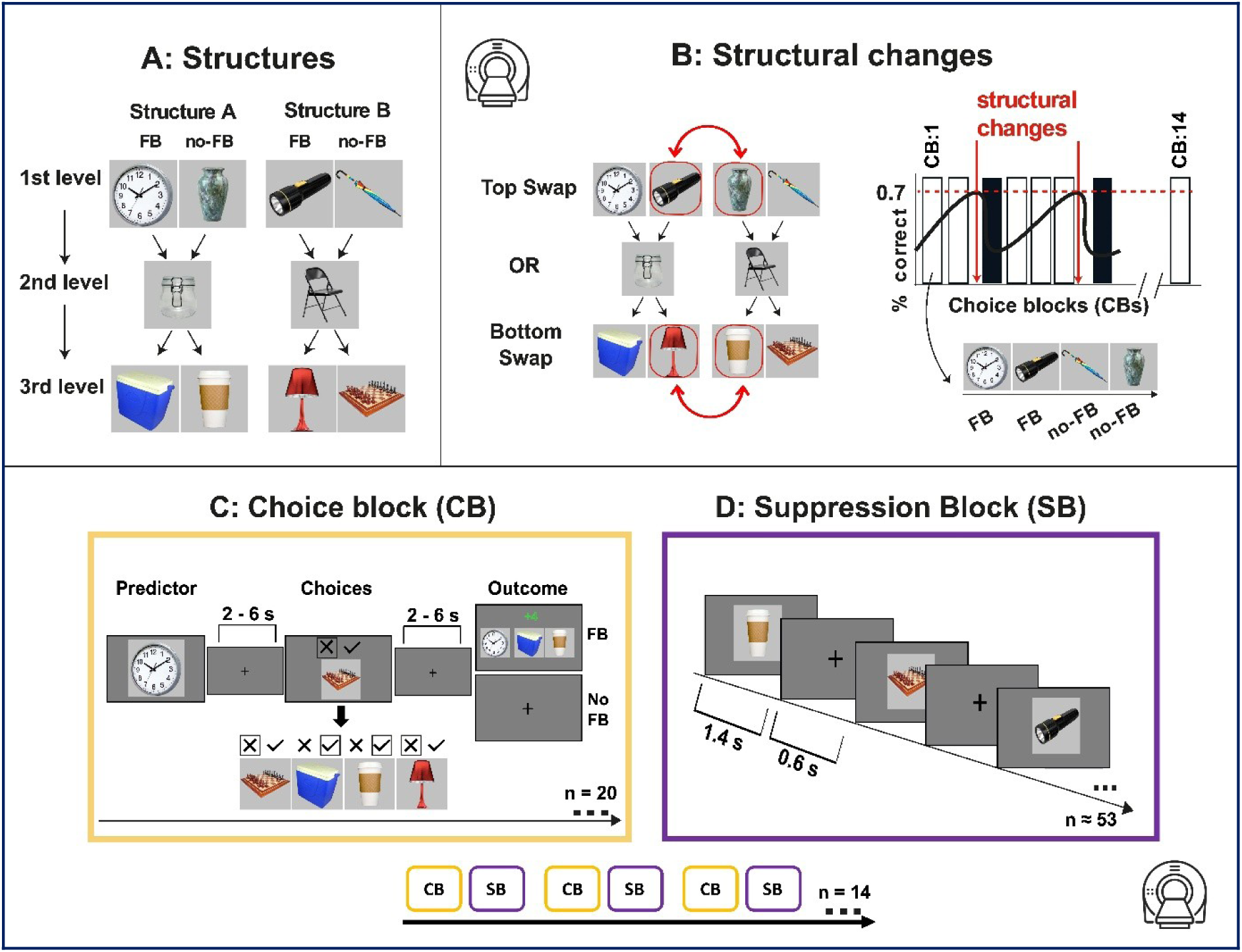
Tracing latent structural changes using a hierarchical structural inference task. **A**. The task required participants to learn two hierarchical structures each comprising three levels of everyday objects. **B**. After this training phase, participants entered the main task inside the scanner, where they were informed that structural changes could occur at either the first or third level of the hierarchy. In each choice block (CB; indicated by white bar) participants had to achieve at least 70% accuracy (black line) before a change could occur (red dotted line) probabilistically (CB following change indicated in black bar; see Methods for details). **C**. During the task, participants selected third-level objects based on whether they were correctly associated with the indicated first-level predictor (e.g. clock) given the current structural relationships. On half the trials they received feedback on the accuracy of their choice (FB), whilst on the other half no feedback was provided (noFB). On FB trials (always presented first in CB), corrective feedback allowed participants to detect that a structural change had taken place; on noFB trials, no such cue was available, and participants had to apply structural inferences drawn from prior FB trials in the same block to guide their choices. Participants were probed about all possible third-level options, with predictors (first-level objects) assigned to the FB or noFB category unbeknownst to the participants. Each choice block (CB; 20 trials, panel **C**) was followed by a repetition-suppression block (SB; approximately 52 trials, panel **D**), in which all possible associative pairs were presented in pseudorandomised order. This design enabled the use of repetition suppression to probe representational patterns (see Figure 3). Each participant completed 14 CB–SB block pairs during the task. Throughout the Results below, all neural analyses were performed on the SB blocks. We refer to the SB block immediately preceding, immediately following, and second-following each structural changepoint as SB-1, SB+1 and SB+2 respectively, in temporal correspondence with the matched behavioural choice blocks (CB-1, CB+1, CB+2).

Participants learned two hierarchical three-level structures, in which two first-level objects were linked to two third-level objects through an implicit middle-layer object. Training was completed prior to scanning over approximately 10 trials of progressive occlusion: the structure was initially shown in full, and on each successive trial more of the structure was hidden, with participants identifying one of the hidden stimuli (Figure 1A). Once participants had learned the structure, they transitioned into the MR scanner. Inside the scanner, participants performed two types of blocks. In choice blocks (CBs; Fig. 1C), they were shown a first-level object and asked to indicate whether a given third-level object was correctly associated with it under the current structure. In interleaved suppression blocks (SBs; Fig. 1D), participants viewed a pseudorandom sequence of approximately 52 stimuli, constructed to ensure that all categories of pairs (see below) were represented while their order was randomised. To maintain attentional engagement during SBs, participants were instructed to detect occasional astronaut targets that appeared at random times in the sequence and to press a button before each target disappeared; missed responses incurred a small monetary penalty deducted from the bonus accumulated in CBs.

To probe flexible adaptation of structural representations, two same-level objects were occasionally covertly swapped between structures (changepoints; Figure 1B), either two first-level or two third-level objects. Participant’s knowledge of these associations was probed through CBs both prior to and following changepoints. For half of the first-level objects, participants received direct feedback on their pairing accuracy (FB; correct / incorrect), whilst for the other half of first-level objects, they received no feedback on their pairing accuracy (noFB; Fig. 1A). Critically, by providing feedback only for some object pairs, the task allowed us to probe whether participants were adequately updating the latent structures of the task and generalizing to objects not provided with feedback. Simpler alternative approaches, such as direct associative (model-free) learning would be insufficient to fully infer on the new structures after each changepoint. Throughout the task, the structure changes were triggered probabilistically based on participants’ performance (unbeknown to participant; see Methods for more details), which allowed us to track how fast participants adapted and generalized following a changepoint. Participants were informed that changes could occur but did not know the timing of such changes.

### Participants use existing knowledge to efficiently adapt and generalize to changes in hierarchical structures

We tested whether participants used their existing knowledge to complete the task and adapt to structural changes. To do this, we measured their performance during choice blocks. We first looked at overall accuracy, calculated as the percentage of correct choices across the entire task. Participants performed well above chance level (50%) (mean = 74.5, sd = 10.6, t(57) = 17.49, p < 0.001; Fig. 2A).

**Figure 2.**
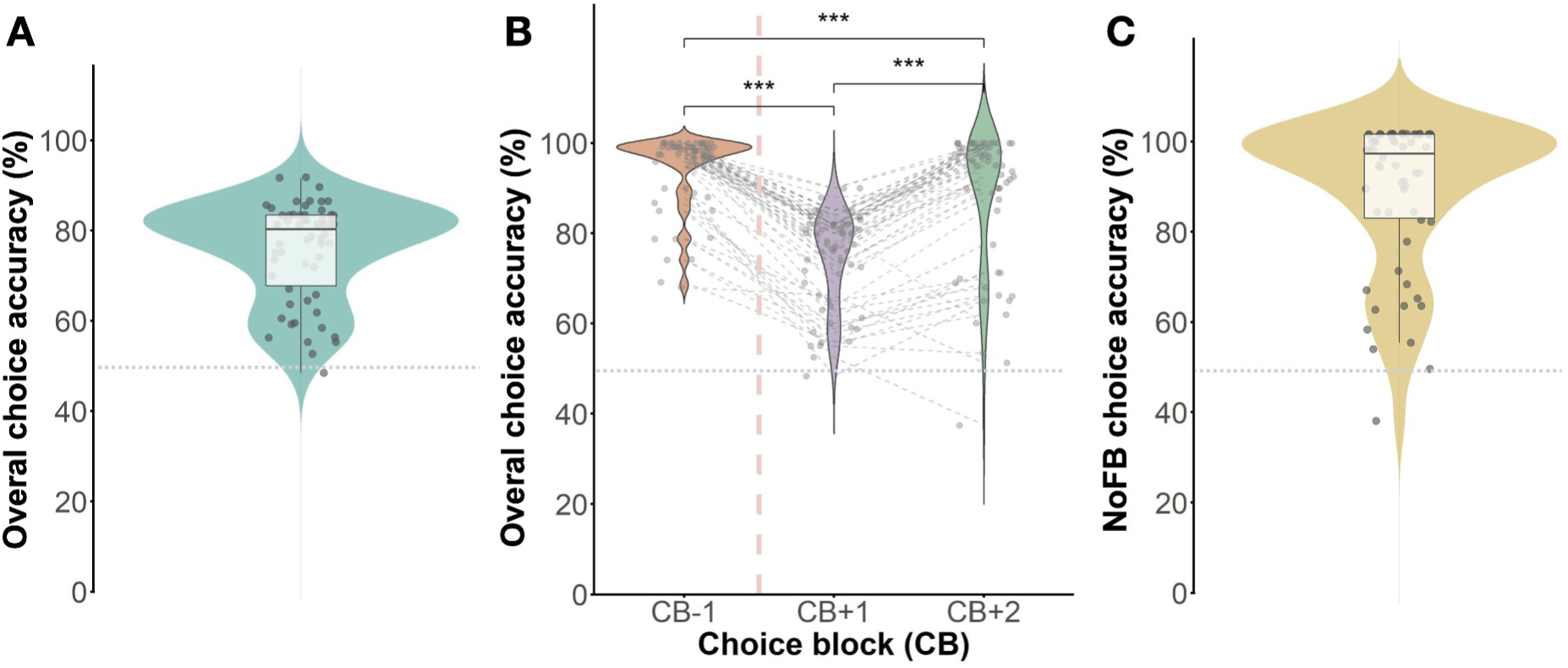
Participants used existing structural knowledge to adapt to changes in the structure of the task. **A.** Participants performed above chance at predicting whether a third-level object was paired or not paired with the displayed first-level object during the choice phase of the task (overall choice accuracy across all trials and third-level objects). **B.** After a structural change, performance initially dropped in the first block (CB+1) but improved at the second block (CB+2; accuracy averaged over 20 trials in each CB). The recovery after structural changes differed substantially between participants (individuals are shown in grey dotted lines). The red dashed line indicates the timing of the changepoint. **C.** Performance in trials without direct feedback remained above chance, illustrating that participants successfully used their structural knowledge to solve this task. Each point in the plot represents a participant’s average performance across all the CBs. **A-C** The dotted horizontal line represents the chance level (50% performance).

To examine how task performance changed over time in response to structural changes, we compared performance in CBs immediately before and after each change. Participants did not know when a structural change would occur, so these timepoints provide the best measure of their adaptability. Performance was highest before the change (CB-1: mean = 94.3, sd = 8.44), dropped sharply in the first block after the change (CB+1: mean = 74.5, sd = 10.7), and partially recovered in the second block (CB+2: mean = 87.8, sd = 15.7). The performance decline from CB-1 to CB+1 and improvement from CB+1 to CB+2 were significant (t(57) = −20.75, p < 0.001; CB+1 to CB+2, t(57) = 11.12, p < 0.001). The rate at which participants recovered after a changepoint varied across individuals, reflecting differences in how effectively they used prior knowledge to adapt to new structures (Figure 2B).

Most importantly, to examine whether participants were able to perform inference using their structural knowledge, we separately analysed choice trials without feedback (noFB; Fig. 1B) and found that participants performed well above chance level (mean = 87.7, sd = 15.9, t = 17.95, df = 57, p < 0.001; Figure 2C). This indicates that participants generalised inferred structural relationships from feedback to no-feedback trials, beyond what could be explained by simple stimulus–response learning. These results highlight their ability to flexibly update and apply structural knowledge.

### Latent structures are represented in the medial OFC

To examine how structural associations were represented and updated in the brain, we recorded fMRI while participants performed the task (see Methods). Choice blocks were interleaved with short suppression blocks (SBs), in which all stimulus pairs from the two hierarchical structures appeared in pseudorandom order. SBs were designed to elicit repetition suppression — the reduction in neural response that occurs when overlapping populations are repeatedly engaged by associated stimuli (Grill-Spector et al., 2006; Barron et al., 2016) — thereby providing an indirect index of representational overlap between pairs. Each SB contained four pair categories defined relative to the most recent changepoint (Figure 3A): stable pairs (associated before and after the change), outdated pairs (associated only before), novel pairs (associated only after), and irrelevant pairs (never associated within the same structure).

**Figure 3.**
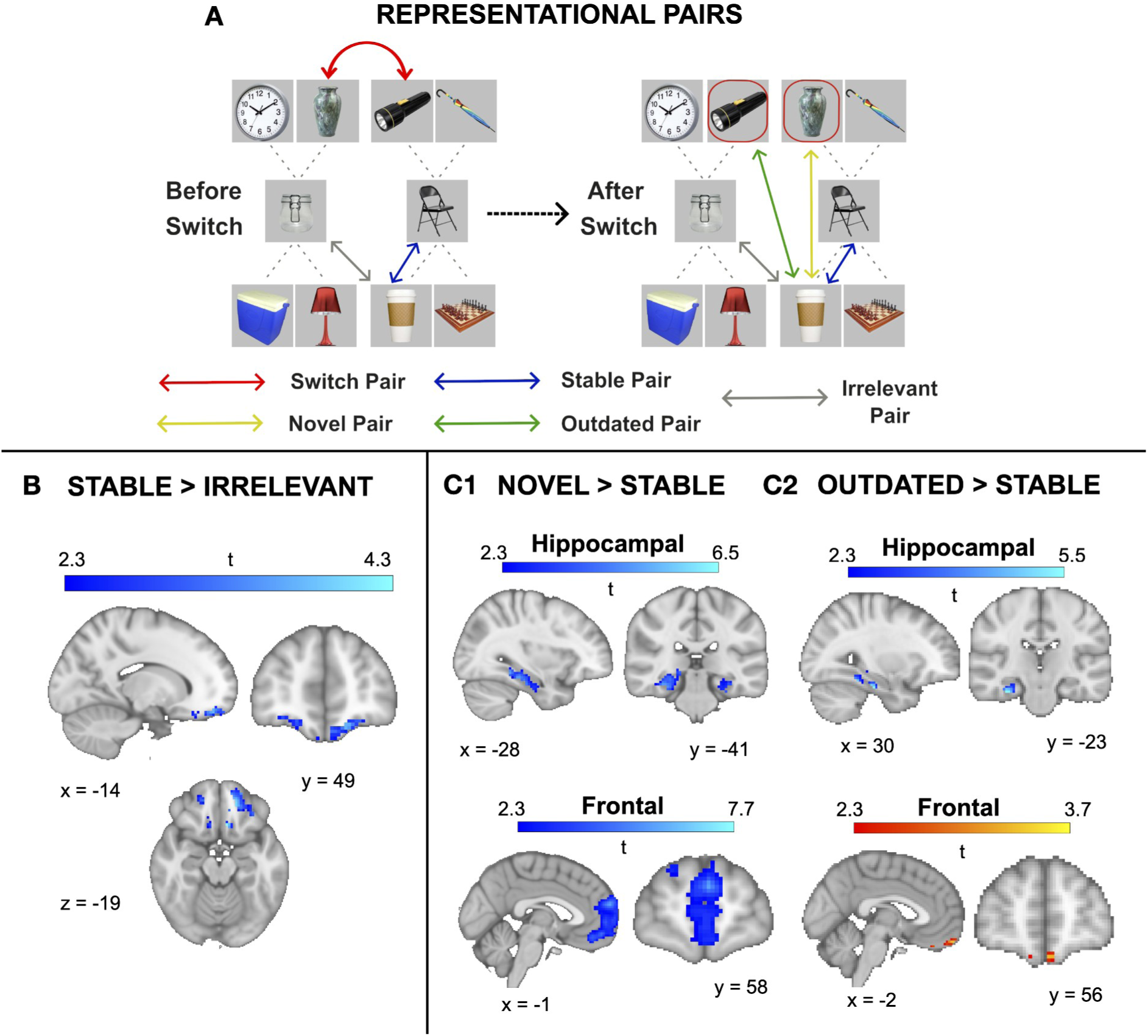
Neural representations of stable, novel, and outdated associative structures across the brain. **A. Categories of representational pairs.** We characterised four types of stimulus–stimulus relationships based on whether pairs belonged to the same latent structure before and after a structural switch. Stable pairs (blue) remained part of the same structure across both phases. Outdated pairs (green) were part of a structure before the switch but no longer associated afterwards. Novel pairs (yellow) were newly formed following the structural change and became part of the updated structure. Irrelevant pairs (grey) were never part of the same structure. Dotted lines illustrate all possible associative links within each structural configuration, while coloured lines highlight example pair types. Note that all possible pairs of each category were included in the analysis at each relevant time point. **B. Stable latent structures are represented in medial orbitofrontal cortex.** Stable pairs showed greater representational overlap relative to irrelevant pairs in medial orbitofrontal cortex (mOFC; MNI: −14, 49, −19; t = 4.3; p < 0.05). This is reflected in reduced BOLD signal for stable relative to irrelevant pairs, consistent with stronger associative encoding for structurally consistent relationships. This effect was observed across all choice blocks, irrespective of whether a structural change had occurred. **C. Persistent representations of novel and outdated pairs across the task (novel > stable; outdated > stable). C1.** Novel pairs, compared with stable pairs, showed greater representational overlap (reduced signal) in bilateral hippocampus (MNI: −28, −41, −13; t = 6.5; p < 0.05) and anterior medial frontal cortex (MNI: −1, 58, 20; t = 7.7; p < 0.05). This reflects the emergence of new associative structures following a switch, specifically distinguishing novel from stable relationships. **C2.** In contrast, outdated pairs showed greater representational overlap relative to stable pairs in a more posterior hippocampal region (MNI: 30, −23, −24; t = 5.5; p < 0.05) and medial orbitofrontal cortex (MNI: −2, 56, −21; t = 3.7; p < 0.05). This pattern indicates residual encoding of previously valid associations despite their removal from the current structure. Specificity vs. irrelevant pairs is shown in Supplementary Figure 4.

Following established RS approaches (Klein-Flügge et al., 2013; Barron et al., 2016; Boorman et al., 2016; Rustichini et al., 2017; see Methods), we focused exclusively on SB blocks for fMRI analyses and contrasted pair categories under the principle that the neural response to the second stimulus of an associated pair is reduced relative to that of an unassociated pair. We applied this logic both across all SBs of the task and within the SBs flanking each changepoint, depending on the analysis below.

We first asked whether mOFC encodes the stable component of a latent structure, as previous work suggests (Wilson et al., 2014; Hunt et al., 2015; Schuck et al., 2016; Klein-Flügge et al., 2019; Boorman et al., 2021). Across all SBs, the stable > irrelevant contrast revealed stronger representational overlap in medial OFC (mOFC; MNI: −14, 49, −19; t = 4.3; p < 0.05, small-volume TFCE-corrected for mOFC ROI, see Methods; Figure 3B). Consistent with the previous literature, stable structural knowledge is therefore represented in mOFC.

### Dynamic representations linger in distinct hippocampal and frontal areas unique for novel and outdated associations

We next asked what happens when latent structural associations change and are replaced by new ones. To address this, we examined neural representational signals across the task, tracking how stimulus relationships evolved over successive structural changepoints (see Methods).

We first focused on newly formed associations. Novel pairs showed greater representational overlap than stable pairs in the bilateral hippocampus (MNI: −28, −41, −13; t = 6.5; p < 0.05, small-volume corrected) and anterior medial frontal cortex (amFC; MNI: −1, 58, 20; t = 7.7; p < 0.05, small-volume corrected; Fig. 3C1), consistent with prior work implicating these regions in abstract and inferred representational learning (Theves et al., 2021; Samborska et al., 2022). Here and below, stronger (i.e. more negative) signals reflect greater representational overlap between stimulus pairs.

We then investigated whether associations that were no longer relevant continued to shape neural representations. Outdated pairs showed greater representational overlap than stable pairs in posterior hippocampus (MNI: 30, −23, −24; t = 5.5; p < 0.05), but reduced overlap in mOFC (MNI: −2, 56, −21; t = 3.7; p < 0.05; Fig. 3C2). This dissociation suggests that although outdated hidden associations continue to be represented in hippocampus, they are selectively underrepresented within mOFC. In other words, once associations become behaviourally irrelevant, they are no longer maintained within the active structural model encoded in mOFC, even though a memory trace of the prior relational structure persists in the hippocampus. Notably, both hippocampal and amFC signals (but not mOFC) remained robust when novel and outdated pairs were compared against irrelevant pairs (Supplementary Fig. 4), indicating that both forms of dynamic representation persist in hippocampus across the task, whereas mOFC preferentially reflects currently stable task structure.

### Novel representations migrate ventrally in the amFC as they consolidate

Having identified distinct persistent neural signals associated with different types of structural representations, we next examined how the novel-pair signal evolved across the timepoints when these structural changes took place. We thus contrasted novel pairs with their corresponding irrelevant pairs at SB+1 and at SB+2 to derive a measure of representational overlap at successive timepoints following the changepoint. At SB+1, when the novel association was likely not yet fully integrated into the existing structure, a large cluster spanning the amFC was observed (MNI: 33, 56, 2; t = 7.7; p < 0.05, small-volume TFCE-corrected; Fig. 4A). By SB+2, when the novel representation was more established (see performance data in Fig. 2), we no longer observed a representational signal in amFC. Instead, the contrast yielded a distinct cluster in mOFC (MNI: −8, 60, −21; t = 5.2; p < 0.05, small-volume TFCE-corrected; Fig. 4B), the same area implicated in stable representations in earlier analyses (Fig. 3B). This pattern illustrates how neural activity evolves as the brain adapts to structural change; shifting from initial encoding of new relationships in amFC to their consolidation and integration into the stable representational system in mOFC, following a dorsal-to-ventral progression along the anterior medial wall.

**Figure 4.**
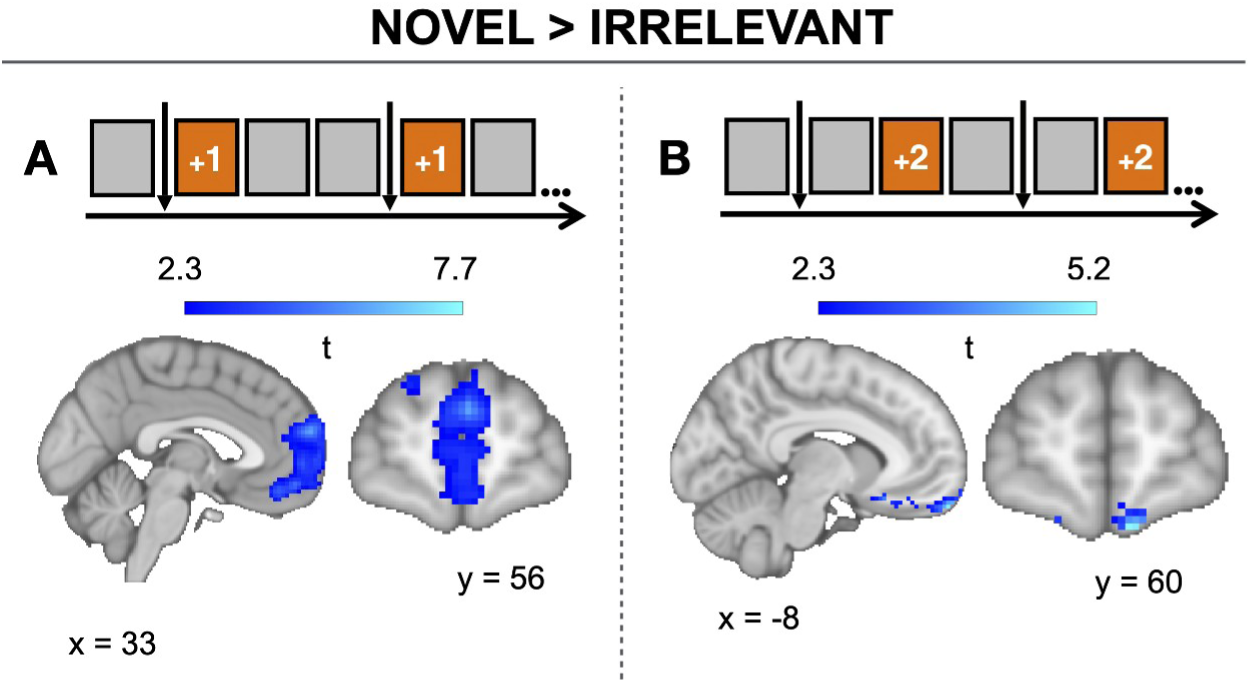
Novel pair representations migrate from amFC to mOFC across consecutive SBs (novel > irrelevant). **A**. Immediately after a changepoint (SB+1), the novel > irrelevant contrast revealed greater representational overlap in amFC (MNI: 33, 56, 2; t = 7.7; p < 0.05). **B**. At the second SB after the change (SB+2), the same contrast yielded a cluster in mOFC (MNI: −8, −32, −21; t = 5.2; p < 0.05), a region linked to stable representations. This temporal shift is consistent with novel representations being initially encoded in a more abstract anterior medial space before being integrated into the stable representational system, paralleling the behavioural recovery observed across the task. The squares at the top indicate each SB, with the vertical arrow marking the changepoint and the horizontal axis the course of the task. Orange squares mark the SB block included in the analysis (SB+1 for A; SB+2 for B).

### Frontopolar and hippocampal regions drive dynamic updating of outdated representations

To understand how formerly relevant associations transition to irrelevant representations, we investigated how representational signal strength changes from before to after a changepoint, i.e. when certain pairs changed their relevance in the task structure (see Methods for further details).

We contrasted previously-relevant, now-outdated pairs before versus after a changepoint (SB+1 vs SB-1). We observed an increase in signal in the bilateral hippocampus (MNI: 19, −8, −16; t = 5.1; p < 0.05, small-volume TFCE-corrected; Fig. 5A) and a reduction in the frontopolar cortex (FP; MNI: 33, 56, 2; t = 4.8; p < 0.05, small-volume TFCE-corrected; Fig. 5B). To visualise the temporal profile of these signals, we extracted parameter estimates from each cluster at SB-1, SB+1, and SB+2 (Fig. 5, right panels). The FP showed a transient peak at SB+1 returning toward baseline by SB+2 (SB+2 compared to SB+1: t(48)=−2.22, p=.031), whereas the hippocampus showed the inverse profile (SB+2 compared to SB+1: t(48)=3.50, p=.002). These patterns are consistent with both regions transiently engaging with outdated pairs at the moment of the changepoint and quietening down once the new structure stabilises. Notably, this dynamic fluctuation closely parallels behavioural adaptation to structural change (see Fig. 2B).

**Figure 5.**
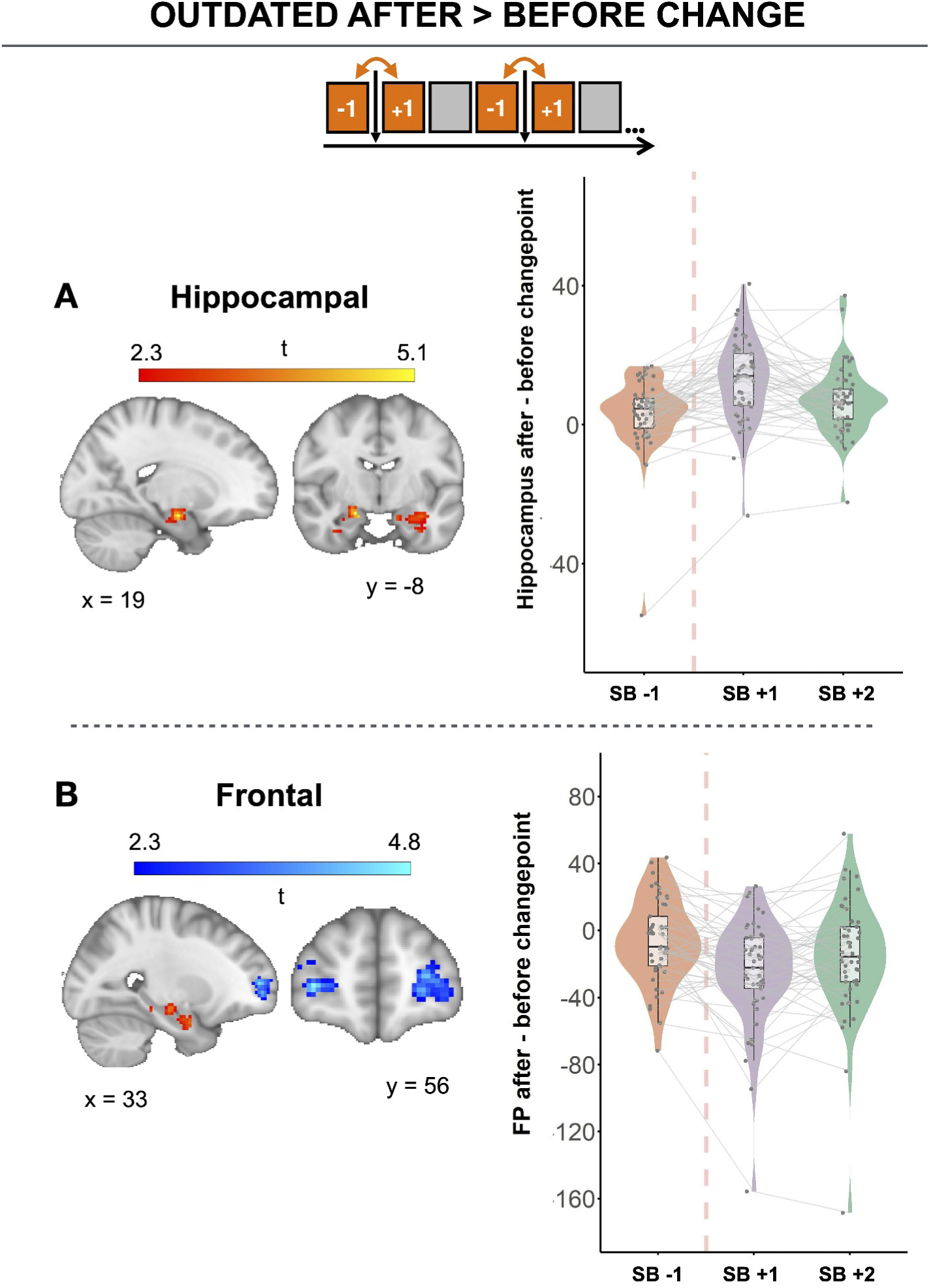
Transient signal modulation in hippocampus and frontopolar cortex for previously-relevant, now-outdated associations (outdated SB+1 > SB-1). **A.** Immediately following the changepoint (SB+1), the anterior hippocampus (MNI: 19, −8, −16; t = 5.1; p < 0.05) showed an elevated signal for outdated pairs relative to SB-1. **B.** Concurrently, the bilateral FP (MNI: 33, 56, 2; t = 4.8; p < 0.05) showed a reduced signal for outdated pairs relative to SB-1. The squares at the bottom indicate each SB across the course of the task, with the vertical arrow marking the changepoint. Orange squares mark the SB blocks contributing to the contrast (SB-1 and SB+1). Right panels display violin plots of parameter estimates extracted from each activation mask at SB-1, SB+1, and SB+2, presented for visualisation only.

### Novel pair processing in amFC and hippocampus tracks individual differences in adaptation

Although all participants were able to learn and adapt to changes in latent structure, there was substantial variability in how rapidly they did so (see Fig. 2B). To examine whether the regions identified above for novel-pair encoding (amFC and anterior hippocampus; Fig. 5) also tracked this individual variability, we split participants into high and low performers based on a median split of accuracy at CB+1 (high performers: mean accuracy = 98.9, sd = 1.34; low performers: mean = 79.9, sd = 12.4) and compared the BOLD response to novel pairs at SB+1 between the two groups.

Comparing novel-pair BOLD response between performance groups at SB+1, we observed greater activation in high than low performers in amFC (MNI: 1, 52, 10; t = 4.3, p < 0.05, small-volume TFCE-corrected) and posterior hippocampus (MNI: −31, −32, −15; t = 5, p < 0.05, small-volume TFCE-corrected; Fig. 6). The same regions implicated above in encoding novel structural relations (Fig. 3B) were thus recruited more strongly in participants who adapted most rapidly to structural change, consistent with prioritised processing of novel structural relations supporting faster behavioural recovery.

**Figure 6.**
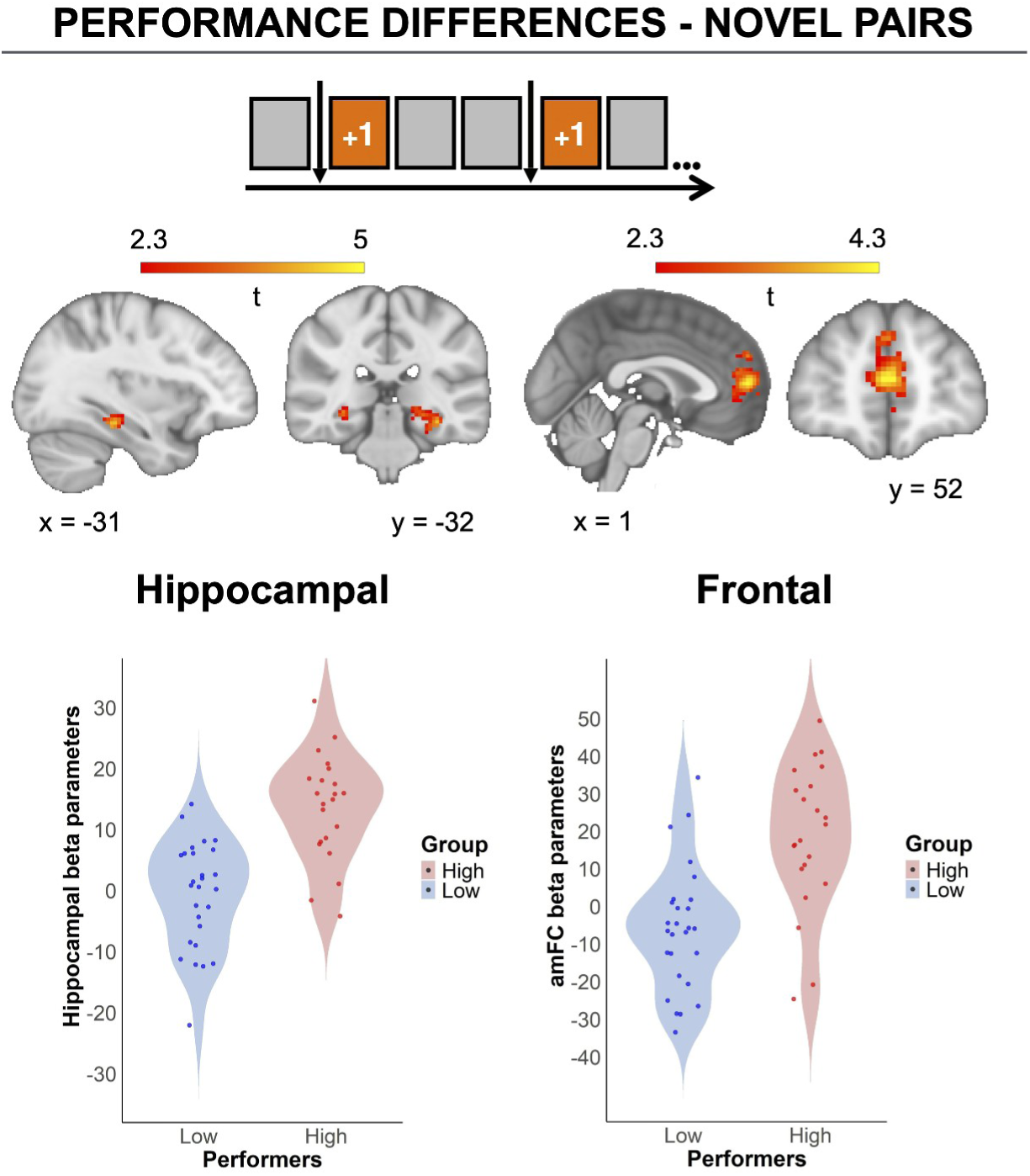
Greater amFC and hippocampal BOLD response to novel pairs in high versus low performers (SB+1). High performers showed elevated BOLD response to novel pairs in amFC (MNI: 1, 52, 10; t = 4.3) and posterior hippocampus (MNI: −31, −32, −15; t = 5; both p < 0.05, small-volume TFCE-corrected). Violin plots display parameter estimates extracted from each cluster for high and low performers (median split) for visualisation. The squares at the bottom indicate that the contrast was computed on the SB block immediately following each structural changepoint (SB+1), temporally aligned with the corresponding choice block (CB+1) at which behavioural recovery is initiated.

## DISCUSSION

Building an internal representation is only adaptive if it accurately reflects the current structure of the environment. This is particularly important in dynamic settings where latent associations can change rapidly and without explicit signalling. Flexible updating of these representations is therefore a prerequisite for cognitive mapping. Here we show that humans update such representations through inference, often in the absence of direct experience for many affected pairs, and that this process engages a fronto-hippocampal network.

In line with prior work, we find that stable relational structures are consistently represented in mOFC, a likely hub for internal cognitive maps, be they spatial or non-spatial, as found across human and non-human animals (Boorman et al., 2016; Klein-Flügge et al., 2019; Bradfield et al., 2018; Bein & Niv, 2025). This pattern suggests that mOFC activity may support the persistence of latent associations, providing a potential foundation for flexible updating when environmental contingencies change. Identifying mOFC as the locus of these enduring representations allowed us to examine how such maps reorganise in response to sudden structural changes.

Our findings show that, despite rapid behavioural adaptation, neural representations of changing structures are separable from stable-structure representationsdistinct and persist beyond the period of behavioural adjustment. Newly formed associations engaged frontal and hippocampal regions, particularly anterior medial frontal cortex (amFC) and anterior hippocampus, with spatially distributed activation patterns. This profile is consistent with a role for these regions in encoding and updating structural change, in line with prior work implicating them in novelty detection and generalisation (Barry et al., 2012; Chen et al., 2020; Bongioanni et al., 2021; Park, S. A. et al., 2021b). One candidate mechanism linking these regions is grid-like coding in entorhinal and prefrontal regions, which provides an abstract coordinate system for relational structures and may drive downstream hippocampal and frontal engagement when structural positions require updating (Park et al., 2020; Garvert et al., 2017; Boorman et al., 2021). Converging MEG evidence further indicates that the rapid construction of hierarchical structure may rely on fast neural sequences that dynamically bind relational roles (Lyu et al., 2025). Whether such grid-like remapping underpins the updating dynamics we observe is an open question for future work.

In contrast, associations that were no longer relevant (outdated pairs) exhibited lingering activity most prominently in a posterior region of hippocampus, consistent with this subregion’s established role in familiarity-based processing and the retention of fine-grained, item-specific memory traces (Strange et al., 1999; Daselaar et al., 2006; Hawco & Lepage, 2014). This posterior localisation stands in direct contrast to the anterior hippocampal engagement observed for novel pairs, and reflects a functionally meaningful gradient along the hippocampal long axis: the anterior hippocampus supports coarse, schema-level representations biased towards pattern completion, whereas the posterior encodes specific associative details biased towards pattern separation (Poppenk et al., 2013; Audrain et al., 2022). The lingering signal for outdated pairs may therefore reflect the retention of precise, item-level representations of previously learned pairings that persist independently of their current behavioural relevance. Such a mechanism, may support transitional periods during structural reconfiguration (Evensmoen et al., 2013; Dandolo & Schwabe, 2018), providing a residual memory scaffold against which new associations can be differentiated and integrated.

Importantly, structural updating appears to involve a transitional processing stage, a kind of representational limbo, before representations being either integrated into a stable schema or discarded. For novel associations specifically, we observe initial recruitment of the amFC, which later shifts ventrally to the mOFC. This temporal gradient is consistent with proposals that anterior orbitomedial regions carry more abstract relational structure, whereas ventral-posterior mOFC encodes more specific, low-dimensional state representations that can support model-based control (Klein-Flügge et al., 2022; Bein & Niv, 2025). To our knowledge, this is the first such characterisation in humans of newly inferred latent structures traversing an intermediate amFC stage before stabilising in mOFC within a single learning episode. In the context of our task — where participants must construct and update a relatively complex relational schema to support rapid generalisation — this redistribution from amFC to mOFC may be particularly important: only once representations are embedded in a stable mOFC map can they flexibly support adaptive choice when the underlying structure changes.

Conversely, outdated representations initially recruit frontopolar cortex (FP), a region consistently implicated in counterfactual evaluation, prospective planning, and cognitive control during rule changes (Boorman et al., 2011; Barbey et al., 2009; Yoshida et al., 2010; Boschin et al., 2015; Nee & D’Esposito, 2016; Badre & Nee, 2018; Miyamoto et al., 2021; Kroger & Kim, 2022; Miyamoto et al., 2024). In our data, outdated pairs show an early expression in FP following the changepoint, followed by more sustained representation in anterior-to-mid hippocampus. This temporal profile is consistent with FP contributing to an initial re-evaluation of the prior structure, potentially supporting counterfactual processing, before these representations are maintained or re-expressed in hippocampus. This suggests a division of labour in which FP responds rapidly to structural change, while hippocampus retains longer-lived traces of prior associations.

The dynamics we observe for novel and outdated pairs sharpen this division of labour between frontal cortex and hippocampus. Around each changepoint, frontal regions (amFC, mOFC, FP) rapidly reconfigure to represent only those pairwise relations that remain structurally or behaviourally relevant for guiding choice, whereas anterior–mid hippocampus continues to encode previously-relevant associations even after they become outdated, both as a sustained signal across the entire task (Fig. 3C) and as a transient response immediately following each changepoint (Fig. 4). Conceptually, this is consistent with the proposal that the hippocampus acts as a flexible, relatively unconstrained processor of associations, whereas mOFC representations are more tightly anchored to immediately biologically significant outcomes (see review by Wikenheiser & Schoenbaum, 2016) and extends this framework to additional frontal regions involved in representing the choice-relevant subset of structural relations. Convergent neural evidence in macaques further supports a distributed fronto-hippocampal circuit underpinning flexible task control following changes in task contingencies (Marché et al., 2026).

Individual differences in adaptation tracked the engagement of these regions for novel associations: participants who recovered most rapidly after a changepoint showed greater BOLD response to novel pairs in amFC and posterior hippocampus immediately after the changepoint, consistent with prioritised processing of new structural relations underpinning faster behavioural adaptation. This finding sits within a broader literature implicating fronto-hippocampal circuitry in the rapid assimilation of new information into existing structural knowledge. Hippocampal object representations have been shown to reorganise dynamically as new conceptual knowledge is acquired (Mack, Love & Preston, 2016), and pre-existing schemas in mPFC accelerate the encoding of compatible new associations, in both rodents (Tse et al., 2007) and humans (van Kesteren et al., 2012). Complementing this, novelty exposure has been shown to reset ventral hippocampal–prefrontal connectivity in rodents, weakening the encoding of an established strategy and enabling rapid acquisition of a new one (Park, A. J. et al., 2021a), whilst in humans, individual differences in mPFC abstract task representations and hippocampal–prefrontal coupling track variability in flexible behaviour and set-shifting performance (Uddin, 2021). Within this framework, our findings suggest that the magnitude of the early novelty response in amFC and hippocampus indexes how rapidly a participant integrates a new structural relation with prior knowledge.

Cognitive inflexibility is a core feature of several psychiatric conditions and is often characterised by impairments in updating internal models in response to changing environmental demands. Within this framework, the individual differences observed here—specifically, variability in the early amFC and hippocampal response to novel structure—may reflect differences in the efficiency with which new relational information is integrated into existing representations. Disruptions to this process could lead to a failure to incorporate new structure, alongside the persistence of outdated associations, a pattern characteristic of rigid cognition observed in disorders such as obsessive–compulsive disorder (Gruner & Pittenger, 2017; Hauser et al., 2017; Robbins, 2022; Uddin, 2021). A natural extension of the present work is therefore to test whether these neural markers predict symptom severity or treatment response in clinical populations.

In summary, our findings characterise the spatiotemporal dynamics of structural learning in the human brain. Stable relational structure is encoded in mOFC, whereas novel and outdated representations recruit broader fronto-hippocampal networks that evolve dynamically over time before being stabilised or attenuated. Individual differences in the engagement of these networks during novel structure processing predict behavioural flexibility, providing insight into the neural mechanisms supporting adaptive cognition. Future work should test whether these neural signatures predict adaptation in patient populations and whether they generalise to other forms of structural learning beyond hierarchical task structures.

## METHODS

### Participants

A total of 51 neurologically and psychiatrically healthy adults participated (age = 25.6 ± 5.6 years, range 18-40; 23 male), all with normal or corrected vision and meeting MRI safety criteria. Participants were drawn from 124 individuals who completed an online performance screen; 89 met inclusion criteria, 61 were invited, and 10 were excluded before analysis (5 inattentive to task structure, 3 technical errors, 2 excessive motion, 1 anatomical anomaly).

Recruitment was through UCL SONA, London-area flyers, and online recruiting platforms. The sample largely comprised UCL undergraduate and master’s students, with no identifiable selection biases. The study received approval from the UCL Research Ethics Committee (14261/002), and all participants gave informed consent.

### Experimental design and procedure

The study comprised an online behavioural session and a subsequent MRI session. In the online phase, participants completed task instructions, comprehension checks, practice rounds, the main structural inference task, and questionnaires. Only participants exceeding a performance threshold on the online task (> 2 structural changes successfully inferred) were invited to the MRI session.

At the MRI visit, participants received a brief refresher on the task instructions and repeated comprehension checks before entering the scanner. They first learned the two initial hierarchical structures, then performed the main fMRI task, divided into two runs with an optional break. Total scanner time was ∼50 min. A main overview of the task is presented bellow with more details in the Supplementary Methods.

#### Task structure

##### Training and initial structure learning

Before the main task, participants were explicitly shown the hierarchical structure linking three stimulus layers (two first-layer “predictors”, a middle layer, and two third-layer “products” per structure). Everyday objects served as stimuli; online and MRI sessions used non-overlapping stimulus sets to minimise perceptual repetition effects.

Participants learned both initial structures via a progressive-occlusion training task. The full structure was first presented, after which successive trials hid a growing subset of stimuli; on each trial, participants identified one of the hidden stimuli from a set of six candidates. Training proceeded over approximately 10 trials per attempt. Participants who reached ≥ 80% accuracy proceeded to the main task, whilst those who did not repeated the training. No participant failed more than once.

##### Main task

The main task examined how participants formed and updated associative beliefs about the hierarchical structures. Each trial consisted of a predictor phase, a sequence of four binary choices, and an outcome phase. At trial onset, a first-layer predictor stimulus was presented, followed by the four third-layer products in random order. For each predictor–product (PP) pair, participants decided whether the two items belonged to the same underlying structure (“accept”) or different structures (“reject”), earning points for correctly identifying the two associated products and rejecting the two non-associated products (see Figure 1D). First-layer predictors were assigned to either feedback (FB) or no-feedback (no-FB) conditions. In FB trials, participants received outcome feedback indicating the two correct products for the current predictor. In no-FB trials, no feedback was provided and a fixation cross was shown instead, requiring participants to infer the correct associations based on previously learned structure. The mapping between predictors and products was deterministic, and the task was arranged such that, by the time no-FB trials occurred, participants had in principle received sufficient information from FB trials to infer the full structure. Performance-based points were converted into a monetary bonus at the end of the experiment.

##### Choice blocks

Trials were organised into unsignalled choice blocks (CBs). Each participant completed 14 CBs, and within each block all feedback and no-feedback trials for both structures were presented once. At the end of each block, one additional trial with the middle-layer node as predictor was included, yielding 20 trials per CB (five predictors × four choices) and providing a block-wise readout of participants’ associative beliefs.

##### Structural changes

Across the task, the underlying mapping between stimuli occasionally changed: at each changepoint, either two first-layer predictors or two third-layer products exchanged their structural assignments (Figure 1C). Participants were informed that such unsignalled changes would occur but not how many or when, and therefore had to infer them from feedback in FB trials. Only one change occurred at a time, and changes were triggered adaptively once accuracy on FB trials exceeded a 70% threshold, ensuring that the current structure had been learned before it was altered. Each participant experienced between one and six structural changes (median = 5; all ≥ 2; Supplementary Figure 2).

##### Suppression Block

At the end of each choice block, a Suppression Block (SB) of approximately 52 stimuli was presented to measure repetition-suppression signal for all possible pairwise associations. During SBs, participants passively viewed a pseudorandom sequence of single stimuli from the two structures, with the order constructed to ensure that all categories of consecutive-stimulus pair (stable, novel, outdated, irrelevant) were represented (see Supplementary Methods for sampling and timing details). To maintain attentional engagement, participants were instructed to detect occasional astronaut targets that appeared at random times in the sequence and to press a button before each target disappeared; missed responses incurred a small monetary penalty deducted from the bonus accumulated in the choice blocks.

### fMRI

Details of MRI data acquisition, preprocessing, and ROI definition are provided in the Supplementary Methods.

### fMRI data analysis

#### General Linear Model Estimation

Two event-related GLMs were estimated in pre-whitened data space to characterise how different types of pairwise representations were expressed across the task and around each changepoint. All regressors were convolved with FSL’s canonical gamma hemodynamic response function and high-pass filtered at 100 s (fast event-related design; Dale, 1999).

We modelled four categories of stimulus pairs presented in the Suppression Blocks: stable (pairs that belonged to the same structure both before and after a changepoint), novel (pairs that became structurally related only after a change), outdated (pairs that were structurally related before but not after a change), and irrelevant (pairs that never belonged to the same structure). For each SB, every consecutive pair was modelled as a single epoch regressor spanning from the onset of the first stimulus to the offset of the second, with separate regressors for stable, novel, outdated, and irrelevant pairs. The contrast between pair-category regressors provided a proxy of the representational overlap between conditions, following the repetition-suppression principle. Pair classifications were re-assigned per SB based on each pair’s relationship to the most recent structural changepoint.

In the *across-task GLM*, we contrasted stable, novel, and outdated pairs against irrelevant pairs to index representational pair signal based on the repetition suppression principle, that is, pairs embedded in a relational structure were expected to show reduced responses relative to structure-irrelevant pairs. In the *switch-focused GLM*, we examined how novel and outdated representations changed around structural changepoints. Outdated pairs were contrasted with irrelevant pairs in the CB before the change (CB–1), immediately after the change (CB+1), and in a later recovery CB (CB+2), and we directly compared outdated representations before versus after the change. For novel pairs, we contrasted novel versus irrelevant pairs at CB+1 and CB+2, when these pairs first became structurally relevant. For subsequent performance-related analyses, beta estimates for novel, outdated, and irrelevant pairs at these CBs were extracted from significant clusters. In the across-task GLM, we contrasted stable, novel, and outdated pairs against each other and against irrelevant pairs (Fig 3) to index representational pair signal: pairs embedded in a relational structure were expected to show reduced responses relative to structure-irrelevant pairs. In the change-focused GLM, we examined how representations evolved across consecutive SBs around each changepoint. For novel pairs, we contrasted novel versus irrelevant pairs at SB+1 and at SB+2 — the SB blocks immediately and second-following each changepoint, when these pairs had only just become structurally relevant (Fig. 4). For outdated pairs, we contrasted previously-relevant-now-outdated pairs at SB+1 versus the same pairs at SB-1 (i.e. before versus after they became outdated; Fig. 5), identifying regions transiently engaged by obsolete associations at the moment of change.

All GLMs additionally included regressors for predictor onset, the choice phase, feedback, Suppression Block onset and offset, and end-of-task/break screens, as well as standard motion and physiological nuisance regressors (framewise displacement, six rigid-body motion parameters, DVARS-based outlier regressors, CSF signal, and physiological regressors; see Supplementary Methods for full specification).

#### Group-level analyses

For each GLM and contrast, the two runs were first combined at the subject level (fixed effects in FEAT) and then entered into mixed-effects group analyses using FSL’s FLAME. Statistical inference used TFCE (Smith & Nichols, 2009) with permutation-based FWE correction (p < 0.05, 5000 permutations). Group-level inference focused on four a priori ROIs (hippocampal formation, mOFC, FP, and amFC), with all second-level and performance-related analyses small-volume corrected within these ROIs (cluster-defining threshold p < 0.001, peak-level FWE p < 0.05). Whole-brain maps are shown for illustration and thresholded as detailed in the Supplementary Methods.

#### Performance related analyses

To examine performance-related differences in novel-pair processing, we used FSL’s FLAME mixed-effects framework with a median-split factor (high versus low performers) based on optimum performance, defined as accuracy relative to the best possible performance given the structural information available on each choice. We then contrasted the BOLD response to novel pairs at SB+1 between groups (high > low; Fig. 6) within the a priori amFC, hippocampus, mOFC and FP ROIs. Optimum performance controls for what was in principle inferable at each timepoint and thus provides a cleaner index of individual differences than raw accuracy. Multiple comparisons were controlled using TFCE.

#### Statistical analysis

To assess effective inference, we tested whether performance in no-feedback trials exceeded chance using one-sample right-tailed t-tests. Behavioural performance changes around structural changepoints (CB-1, CB+1, CB+2) were examined with paired t-tests. For visualisation of fMRI effects, beta estimates were extracted from significant clusters at SB-1, SB+1 and SB+2; because these clusters were defined by contrasts involving the same SB blocks, the extracted estimates are used for descriptive illustration only and were not subjected to further inferential testing. Where multiple comparisons were performed within an analysis family, p-values were adjusted using Bonferroni correction. Behavioural analyses were conducted in R.

#### Data availability

All behavioural and processed neuroimaging data supporting the findings of this study, including unthresholded statistical maps, are publicly available via NeuroVault at https://identifiers.org/neurovault.collection:23781. Raw neuroimaging data are available from the corresponding author upon reasonable request, subject to ethical approval.

#### Code availability

All original analysis code has been deposited at GitHub and is publicly available as of the date of publication. The behavioural analysis pipeline is available at https://github.com/gtertikas/structural-inference-behaviour and the ROI extraction and visualisation pipeline is available at https://github.com/gtertikas/structural-inference-roi.

## AUTHOR CONTRIBUTIONS

Georgios Tertikas (G.T.) and Nadescha Trudel (N.T.) contributed equally to this work. N.T., Miriam Klein-Flügge (M.K.-F.) and Tobias U. Hauser (T.U.H.) conceived the study and developed the overall research question and design. G.T., N.T., T.U.H. and M.K.-F. designed the methodology and analysis approach, and G.T. implemented the experimental task and analysis code. Data were collected by G.T. and G.T. performed the formal analyses and data curation. G.T. prepared the figures and visualizations. G.T. wrote the original draft of the manuscript, and all authors (G.T., N.T., M.K.-F. and T.U.H.) contributed to reviewing and editing the final version. T.U.H. provided supervision and secured funding and other resources for the project.

## ACKNOWLEDGMENTS

TUH is supported by a Wellcome Sir Henry Dale Fellowship (211155/Z/18/Z) and other project funding from the Wellcome Trust (316955/Z/24/Z). This work was supported by the Carl-Zeiss-Stiftung and the Alexander von Humboldt foundation, precisely the Alexander-von-Humboldt-Professorship award to Peter Dayan. The Max Planck UCL Centre is a joint initiative supported by UCL and the Max Planck Society. The Wellcome Centre for Human Neuroimaging is supported by core funding from the Wellcome Trust (203147/Z/16/Z). MCKF was funded by a Wellcome Trust Henry Dale Fellowship (223263/Z/21/Z), a UKRI-converted ERC Starting Grant (EP/X021815/1) and a Leverhulme Award in Psychology. LLM tools have been used for language editing of this manuscript.

## SUPPLEMENTARY MATERIAL

### Supplementary methods

#### Imaging Behavioural task – further details

##### Training and initial structure learning

At the beginning of the MRI visit, participants were reminded of the task using a shortened version of the instructions from the online session. They then completed a set of multiple-choice comprehension questions designed to probe: (i) understanding of the three-layer hierarchical structure, (ii) the mapping between the first, middle, and third layers with regards to feedback, (iii) the nature of unsignalled structural changes (which layers could change and that only one change would occur at a time). All questions had to be answered correctly before participants could proceed; any incorrect answer led to repetition of the entire question set until all responses were correct.

During the instruction phase, participants viewed explicit diagrams of the two hierarchical structures. Each structure comprised two first-layer “predictors” connected to a single middle-layer node, which in turn connected to two third-layer “products”. Everyday objects were used as stimuli. To minimise carry-over and avoid contamination of repetition-suppression effects, the set of objects used in the MRI session was entirely distinct from that used in the online screening session and every structure was randomly generated and was different per participant. Participants learned the mapping between stimuli and structural positions using a task that resembled a memory card game. On each trial, a partially completed hierarchical diagram was displayed on the left side of the screen, with one or more items missing. On the right, six candidate items were shown. Participants selected the item(s) they believed correctly completed the structure. Feedback indicated whether the chosen items were correct, allowing participants to gradually learn both hierarchical mappings. Training continued until participants reached at least 80% accuracy across trials. If this criterion was not met, the structure-learning block was repeated. No participant required more than one repetition to reach criterion.

##### Main task

As described in the main Methods, each trial comprised a predictor phase, four binary predictor–product (PP) choices, and an outcome phase, with predictors assigned to FB or no-FB conditions. The mapping between predictors and products was deterministic for a given structural configuration: each predictor was linked to exactly two products and never to the remaining two.

The trial schedule was arranged such that no-FB predictors only appeared after participants had, in principle, received sufficient FB evidence to infer the entire mapping between predictors, middle-layer nodes, and products. Accurate performance on no-FB trials therefore required participants to maintain and exploit a coherent relational schema rather than rely on local reward history.

Because each predictor belonged to exactly one of the two structures, products from the other structure were effectively anti-correlated choices: accepting a product from one structure implied that products from the alternative structure should be rejected for that predictor. This property meant that choices reflected both direct associative knowledge and inference over unobserved, structurally implied relationships.

Participants accumulated points for correctly accepting associated products and correctly rejecting non-associated products. Total points across the session were linearly converted into a monetary bonus added to the base payment.

##### Choice blocks

Trials were organised into unsignalled choice-blocks (CBs). Within each CB, the order of first-layer predictors was randomised within feedback condition (FB vs no-FB), while the sequence of FB and no-FB trials across CBs was held constant across participants to maintain a consistent global structure.

The additional middle-layer trial at the end of each CB used the middle-layer node as the predictor. On this trial, participants judged four associations involving two first-layer and two third-layer items (one from each structure), again via binary “accept/reject” responses and without feedback. These middle-layer trials provided an extra readout of cross-structure relational knowledge.

To decorrelate BOLD responses associated with different events, jittered intervals were inserted before predictor onset, between the choice phase and the outcome, and before the next trial. Jitter durations were sampled from a gamma distribution with a minimum of 2 s, a maximum of 6 s, and a mean of approximately 4 s.

##### Structural changes

Structural changes were implemented as swaps of stimulus assignments between the two structures, either at the first layer (swapping two predictors) or at the third layer (swapping two products). Changes were constrained so that either (i) two FB predictors were swapped, or (ii) one FB and one no-FB predictor were swapped; two no-FB predictors were never swapped. This ensured that changes always involved at least one stimulus for which feedback was available, guaranteeing that the new structure was, in principle, inferable.

The set of possible swaps (which specific stimuli could exchange positions) was fixed across participants. However, whether and when each potential change occurred depended on individual performance. Within each CB, if accuracy on FB trials exceeded 70%, a structural change became eligible. To reduce predictability, a change was only implemented with 50% probability when the criterion was met; if no change was implemented, the next time the criterion was exceeded a change was forced. After each change, at least two full CBs elapsed before another change could occur, preventing rapid successive changes.

At the start of the experiment, a structural change was permitted after the first CB because both structures had been explicitly learned beforehand. Across the MRI session, participants experienced between one and six structural changes, with the total number bounded by the overall trial count. All participants experienced at least two changes (median = 5; see Supplementary Figure 2 for the distribution and timing of changepoints). The sequence of structural changes in the MRI task differed from that in the online screening task to minimise expectations about when changes would occur.

##### Suppression Block

At the end of each CB, a Suppression Block (SB) was presented. Each SB began and ended with a green fixation cross. Within an SB, all unordered pairs of the ten stimuli (10 choose 2 = 45 unique pairs) were presented in a continuous sequence. Each pair was shown as two successive images: the first stimulus of the pair, followed by the second stimulus, each separated by brief fixation intervals. The sequence of pairs was pseudo-randomised with the constraint that the second stimulus of one pair could serve as the first stimulus of the next, yielding on average ∼53 images per SB.

Each image was displayed for 1.4 s, followed by a fixation cross for 0.6 s. On every SB, an astronaut icon appeared superimposed on the images on 10 trials, never earlier than 0.5 s after picture onset. Participants were instructed to press the middle button as quickly as possible whenever they saw the astronaut. These catch trials served as an attention check and contributed to the performance-based bonus. Participants were informed that poor performance during SBs could reduce their overall bonus, to promote sustained engagement.

Across CBs and participants, the order and sampling of specific stimulus pairs were re-generated to avoid trivial repetition of identical pair sequences and to decorrelate pair identity from CB number.

#### MRI data acquisition and preprocessing

Visual stimuli were projected onto a screen via a computer monitor. Participants indicated their choices using an MRI-compatible button box. MRI data were acquired using a 64-channel head coil on a 3T Siemens MAGNETOM Prisma Fit system (Siemens). fMRI scans were acquired in axial orientation using T2*-weighted gradient-echo echo-planar imaging (GE-EPI) with multiband acceleration, sensitive to BOLD contrast. Echo-planar imaging (EPI) with sampling after multiband excitation achieves temporal resolution in the subsecond range while maintaining good slice coverage and spatial resolution.

We collected 60 transverse slices of 2.4 mm thickness with an in-plane resolution of 2.4 × 2.4 mm, a multiband acceleration factor of two, a repetition time of 1.5 s, and an echo time of 20 ms. Slices were acquired in an interleaved order with a flip angle of 65° and a phase encoding direction of anterior to posterior. The GRAPPA parallel imaging method was applied with an acceleration factor of two. The acquisition field of view was 216 × 216 mm with a matrix size of 90 × 90. The first five volumes of each block were discarded to allow for scanner equilibration. The selection of the specific scanning sequence settings (ie. TR, TE and multiband factor) were based on an in lab analysis, so at to have the least signal drop in the orbitofrontal cortex, given that it is an a priori region of interest.

High-resolution anatomical images were acquired using a T1-weighted magnetization-prepared rapid gradient-echo (MPRAGE) sequence. Scans were obtained in the sagittal plane with a field of view of 256 × 256 × 176 mm and an isotropic voxel size of 1 mm³. The acquisition parameters included a repetition time (TR) of 2530 ms, an inversion time (TI) of 1100 ms.

Additionally, a field map was acquired using a dual-echo gradient-recalled echo (GRE) sequence in the transverse plane to estimate B0 field inhomogeneities. The field of view was 192 × 192 × 191 mm with a voxel size of 3.0 × 3.0 × 2.0 mm. A total of 64 slices were acquired in an interleaved order with a slice thickness of 2 mm. The repetition time (TR) was 1020 ms, with two echo times (TE1 = 10.00 ms, TE2 = 12.46 ms). The flip angle was set to 90°, and the phase encoding direction was right to left (R >> L).

In addition, physiological recordings were taken during the functional MRI blocks to measure the participant’s pulse and breathing.

#### Data preprocessing

Results included in this manuscript come from preprocessing performed using fMRIPrep 23.2.1 (Esteban et al. (2019); Esteban et al. (2018); RRID:SCR_016216), which is based on Nipype 1.8.6 (K. Gorgolewski et al. (2011); K. J. Gorgolewski et al. (2018)).

##### Preprocessing of B0 inhomogeneity mappings

A B0 nonuniformity map (or fieldmap) was estimated from the phase-drift map(s) measure with two consecutive GRE (gradient-recalled echo) acquisitions. The corresponding phase-map(s) were phase-unwrapped with prelude (FSL; Jenkinson, 2003).

##### Anatomical data preprocessing

A total of 1 T1-weighted (T1w) images were found within the input BIDS dataset. The T1w image was corrected for intensity non-uniformity (INU) with N4BiasFieldCorrection (Tustison et al. 2010), distributed with ANTs 2.5.0 (Avants et al. 2008, RRID:SCR_004757), and used as T1w-reference throughout the workflow. The T1w-reference was then skull-stripped with a Nipype implementation of the antsBrainExtraction.sh workflow (from ANTs), using OASIS30ANTs as target template. Brain tissue segmentation of cerebrospinal fluid (CSF), white-matter (WM) and gray-matter (GM) was performed on the brain-extracted T1w using fast (FSL (version unknown), RRID:SCR_002823, Zhang, Brady, and Smith 2001). Volume-based spatial normalization to two standard spaces (MNI152NLin6Asym, MNI152NLin2009cAsym) was performed through nonlinear registration with antsRegistration (ANTs 2.5.0), using brain-extracted versions of both T1w reference and the T1w template. The following templates were were selected for spatial normalization and accessed with TemplateFlow (23.1.0, Ciric et al. 2022): FSL’s MNI ICBM 152 non-linear 6th Generation Asymmetric Average Brain Stereotaxic Registration Model [Evans et al. (2012), RRID:SCR_002823; TemplateFlow ID: MNI152NLin6Asym], ICBM 152 Nonlinear Asymmetrical template version 2009c [Fonov et al. (2009), RRID:SCR_008796; TemplateFlow ID: MNI152NLin2009cAsym].

##### Functional data preprocessing

For each of the 2 BOLD runs found per subject (across all sessions), the following preprocessing was performed. First, a reference volume was generated, using a custom methodology of fMRIPrep, for use in head motion correction. Head-motion parameters with respect to the BOLD reference (transformation matrices, and six corresponding rotation and translation parameters) are estimated before any spatiotemporal filtering using mcflirt (FSL, Jenkinson et al. 2002). The estimated fieldmap was then aligned with rigid-registration to the target EPI (echo-planar imaging) reference run. The field coefficients were mapped on to the reference EPI using the transform. The BOLD reference was then co-registered to the T1w reference using mri_coreg (FreeSurfer) followed by flirt (FSL, Jenkinson and Smith 2001) with the boundary-based registration (Greve and Fischl 2009) cost-function. Co-registration was configured with six degrees of freedom. Several confounding time-series were calculated based on the preprocessed BOLD: DVARS and region-wise global signals. DVARS was calculated for each functional run, using its implementations in Nipype (following the definitions by Power et al. 2014). The global signal was extracted within the CSF. The head-motion estimates calculated in the correction step were also placed within the corresponding confounds file. Frames that exceeded a threshold of 1.5 standardized DVARS were annotated as motion outliers.

##### ROI analysis

For all the analyses, we report all results within the hippocampal formation, orbitofrontal cortex (OFC), frontopolar cortex (FP), and anterior medial frontal cortex (amFC), as these were our predefined regions of interest (ROIs). These results were identified using an uncorrected cluster-defining threshold of *p* < 0.001, combined with peak-level FWE small-volume correction at *p* < 0.05. ROI masks were derived from Neuroquery based on existing literature. Activation in other regions was considered significant only at *p* < 0.001 (uncorrected), with peak-level FWE small-volume correction applied at the cluster level (*p* < 0.05).

While multiple comparisons within our ROIs were controlled using small-volume correction, all statistical parametric maps presented in the manuscript are unmasked and thresholded at *p* < 0.01 for visualization.

##### Noise regressors

Because of the sensitivity of the blood oxygen level-dependent signal to motion and physiological noise, all GLMs included frame-wise displacement, six rigid-body motion parameters (three translations and three rotation), one outlier regressor based on DVARS and one CSF regressor. Additionally, to remove variance accounted for by cardiac and respiratory responses, a physiological noise model was constructed using an in-house developed Matlab toolbox (Hutton et al., 2011). Models for cardiac and respiratory phase and their aliased harmonics were based on RETROICOR (Glover et al., 2000). The model for changes in respiratory volume was based on Birn et al. (2006). This resulted in 17 physiological regressors in total: 10 for cardiac phase, six for respiratory phase, and one for respiratory volume.

### Supplementary figures

**Supplementary Fig. 1.**
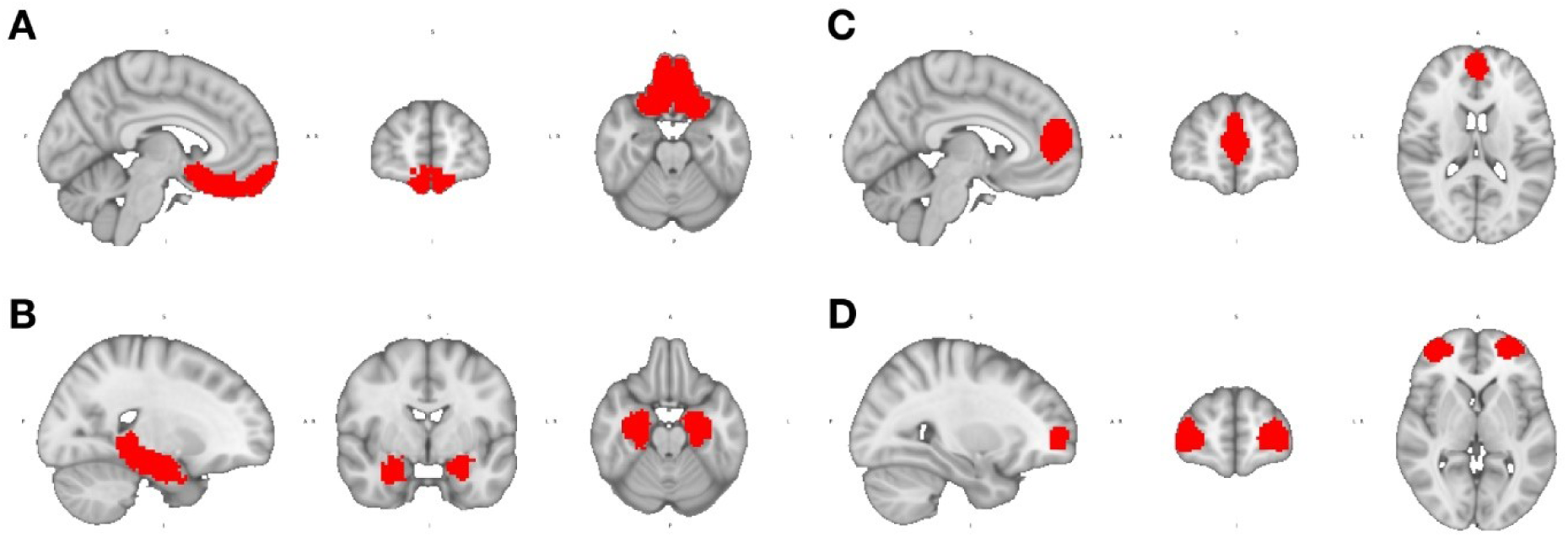
Masks used for the TFCE analysis. **A.** Orbitofrontal cortex (OFC) **B.** Hippocampal area **C.** Anterior medial frontal cortex (amFC) **D.** Frontopolar cortex (FP).

**Supplementary Fig. 2.**
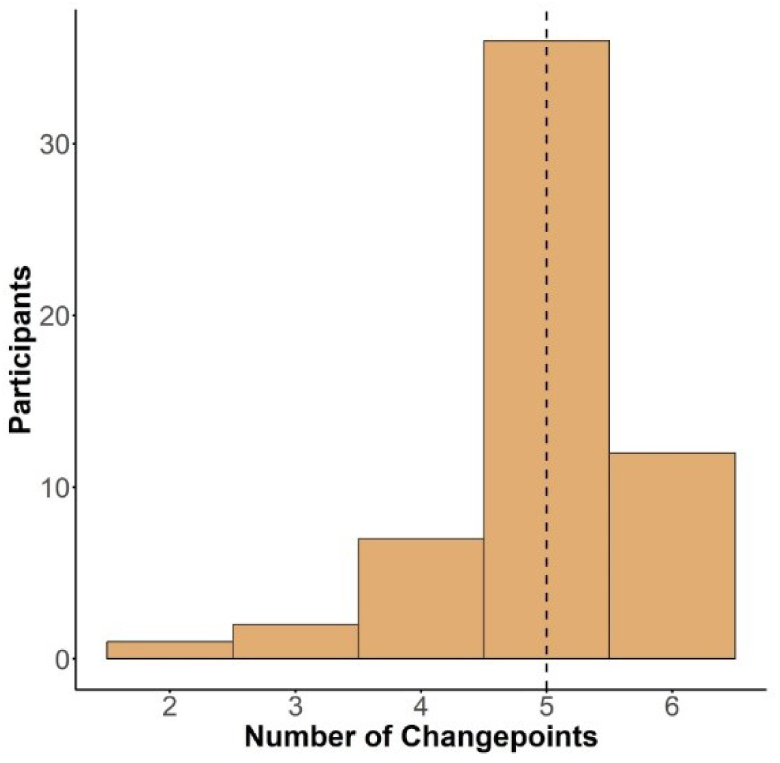
Number of changepoints elicited by participants during the task. Bar plot illustrating the number of changepoint per participant. Dotted line represents the median.

**Supplementary Fig. 3.**
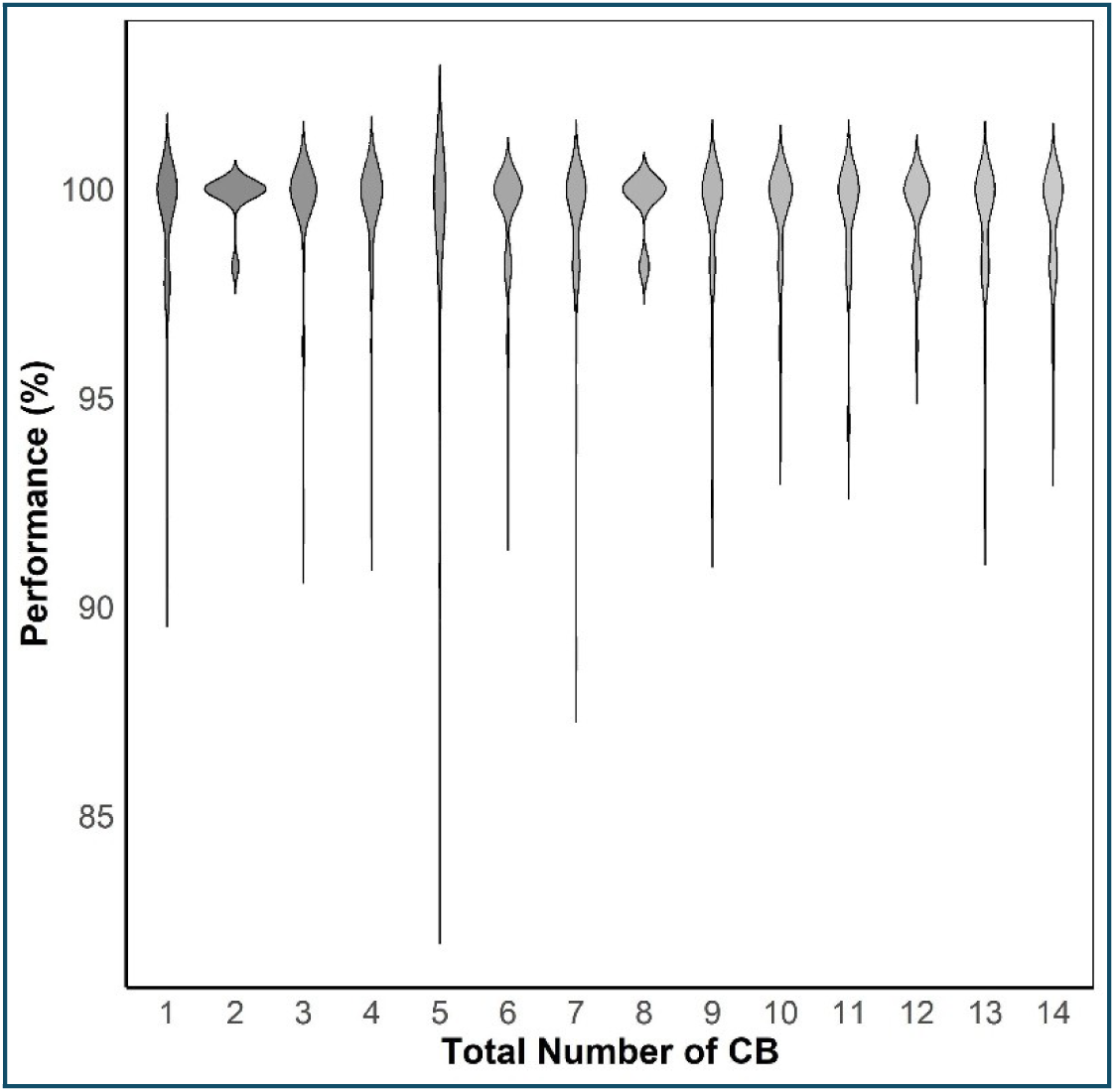
Performance in each Suppression block (SB). Violin plots depicting average performance in the astronaut game during the suppression block (SB) for each choice block (CB). Performance above 85% indicates that participants effectively attended to SB stimuli, allowing for the reliable probing of repetition suppression representations.

**Supplementary Fig. 4.**
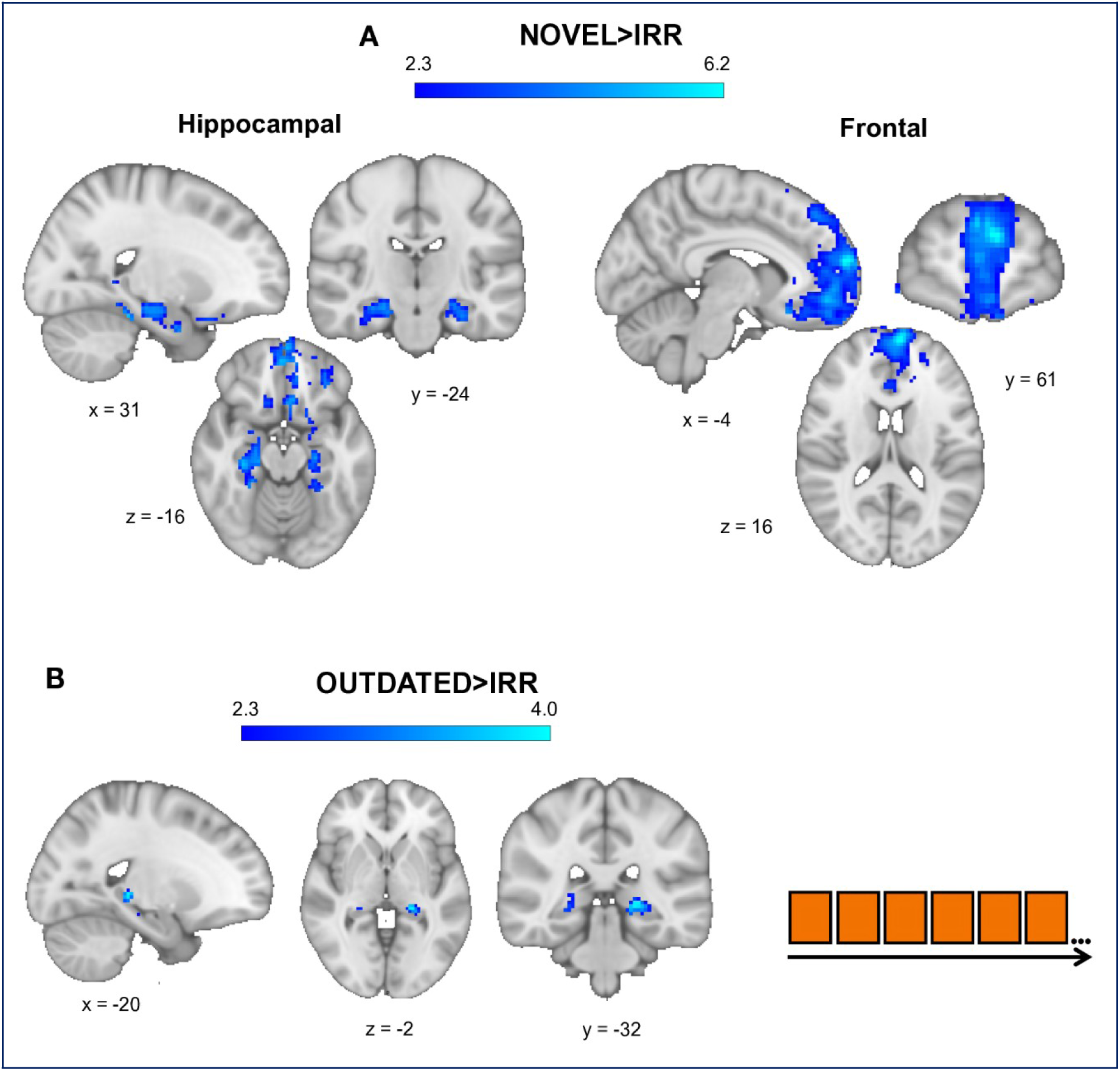
Novel and outdated representational signal compared to irrelevant pairs across time. **A–B.** Distinct neural signals for novel and outdated representation pairs when comparing with irrelevant pairs throughout the task. **A.** Novel representational pairs compared to irrelevant pairs elicited a reduced signal (increased associative signal) in the bilateral hippocampus (MNI: 31, −24, −16; t = 5; p < 0.05) and the anterior medial frontal cortex (amFC) (MNI: −4, 61, 16; t = 6.2; p < 0.05). **B.** In contrast, outdated representational pairs compared to irrelevant pairs had an increased associative signal (reduced neural activity) in a more posterior region of the hippocampus (MNI: −20, −32, −2; t = 4; p < 0.05). Panels A–B are thresholded at *p* < 0.01 (uncorrected) for visualization within the ROIs.

**Supplementary Fig. 5.**
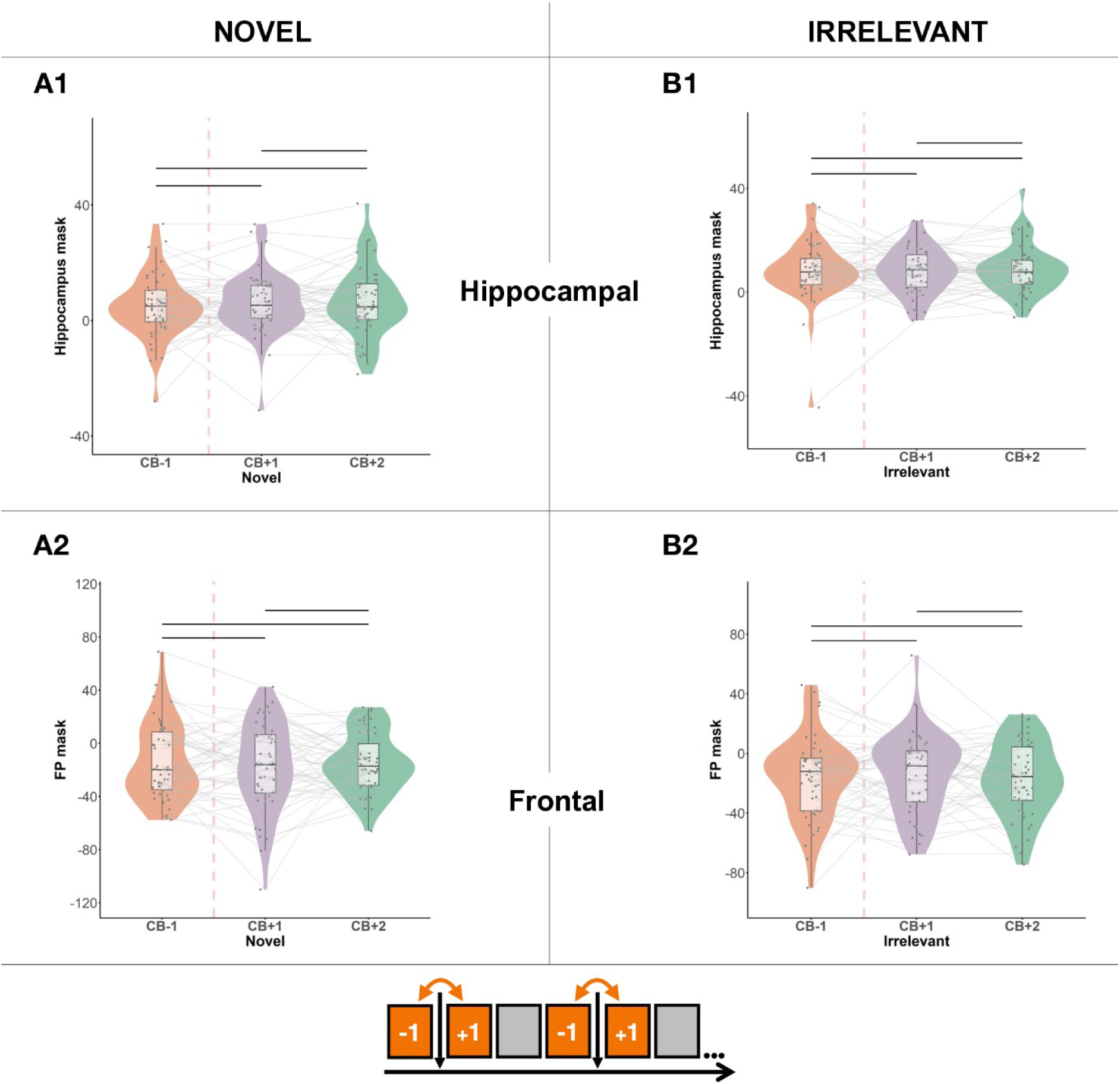
Beta parameter extraction of novel and irrelevant representational signal. The parameters were extracted from the hippocampal and frontal masks of activation of the outdated representation before and after each changepoint. Paired t-tests comparing beta parameters pre- and post-changepoint revealed no statistically significant differences.

**Supplementary Fig. 6.**
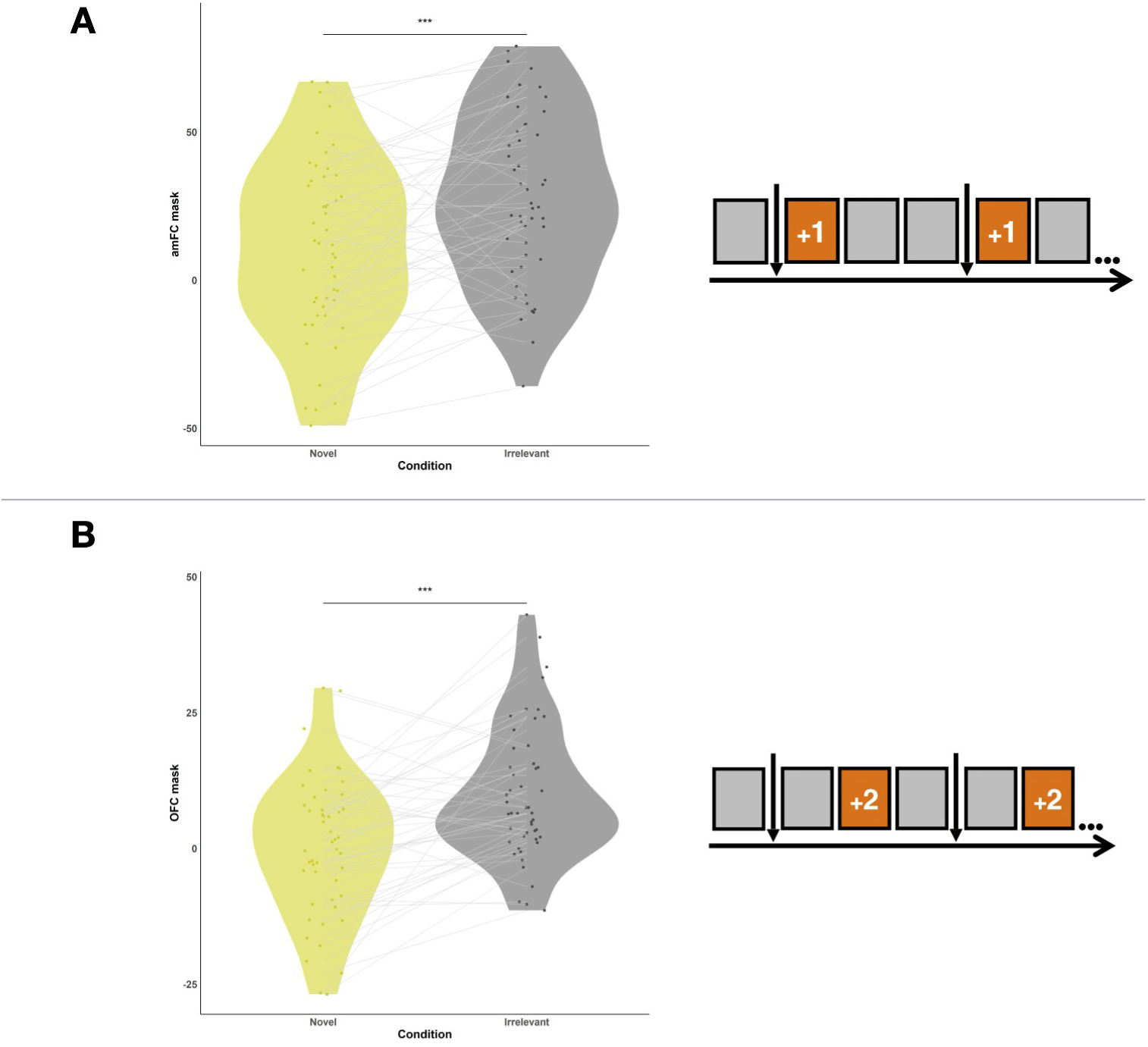
Beta parameters extracted from statistically significant representational contrasts at time points after the changepoint. Violin plots of beta parameters across different representational pairs. Paired t-tests were conducted to assess expected significance. **A.** Novel and irrelevant representational beta activation from the amFC mask at CB+1. **B.** Novel and irrelevant representational beta activation from the mOFC mask at CB+2. Across **A–B**, all beta parameter distributions are equal, suggesting no impact of possible trial inequality between representation types. ***P < 0.001.

