## Supplemental material for "Frontotemporal cortex flexibly adapts latent structural representations"

Supplementary Material for: Frontotemporal cortex flexibly adapts latent structural representations

### SUPPLEMENTARY MATERIAL

### Supplementary methods

#### Imaging Behavioural task – further details

###### Training and initial structure learning

### Supplementary figures

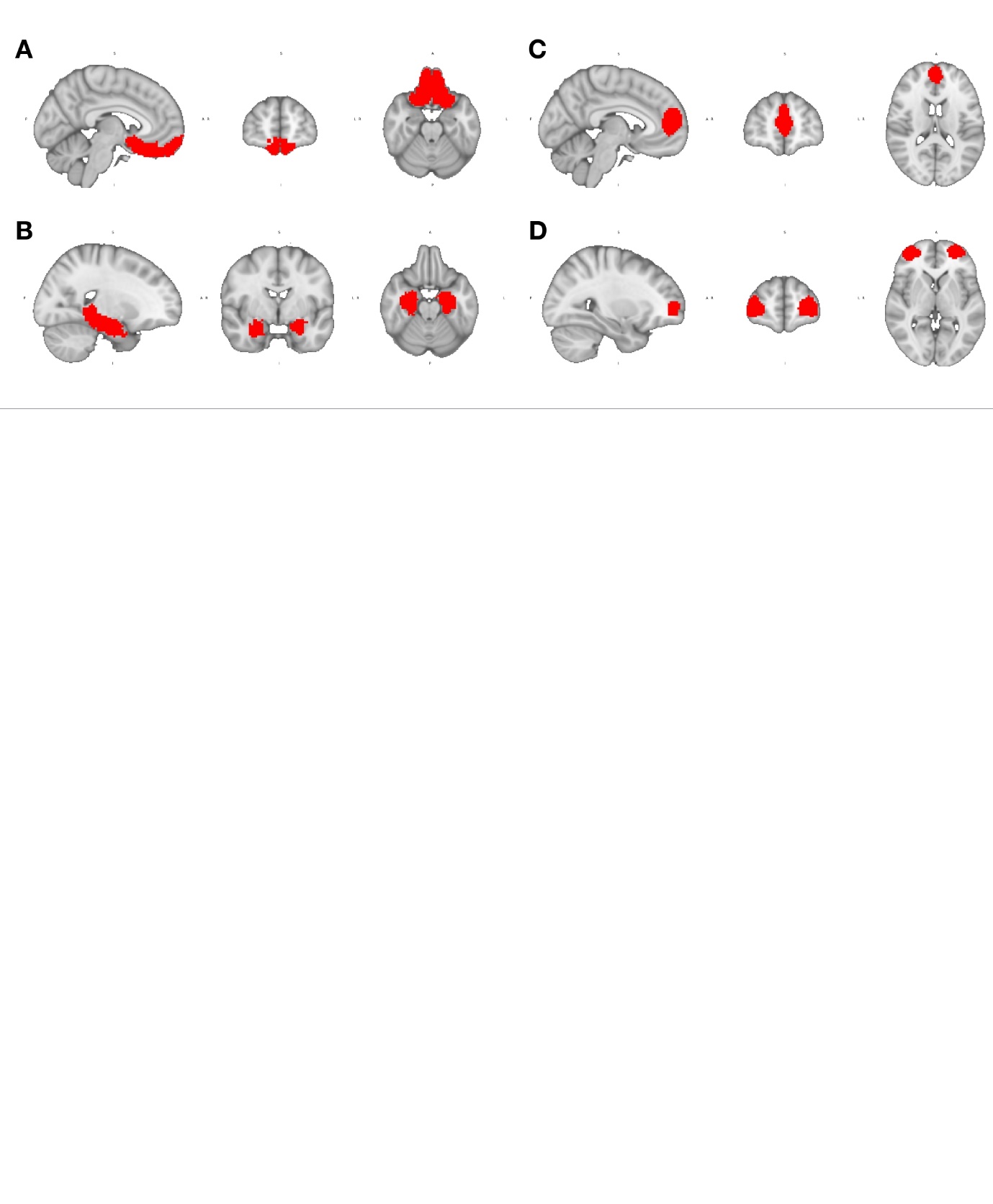

Supplementary Fig. 1 | Masks used for the TFCE analysis.

A. Orbitofrontal cortex (OFC) B. Hippocampal area C. Anterior medial frontal cortex (amFC) D. Frontopolar cortex (FP).

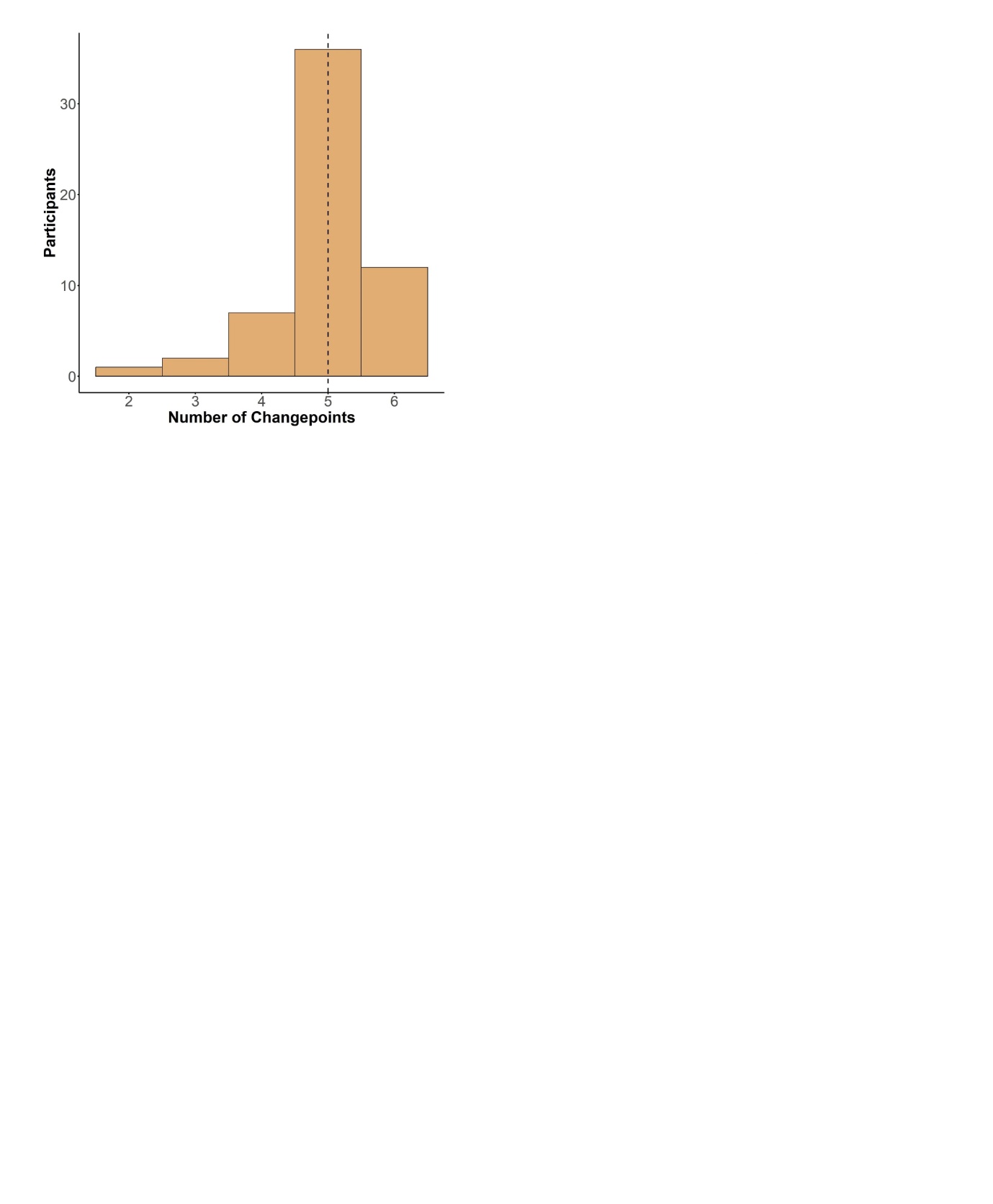

Supplementary Fig. 2 | Number of changepoints elicited by participants during the task.

Bar plot illustrating the number of changepoint per participant. Dotted line represents the median.

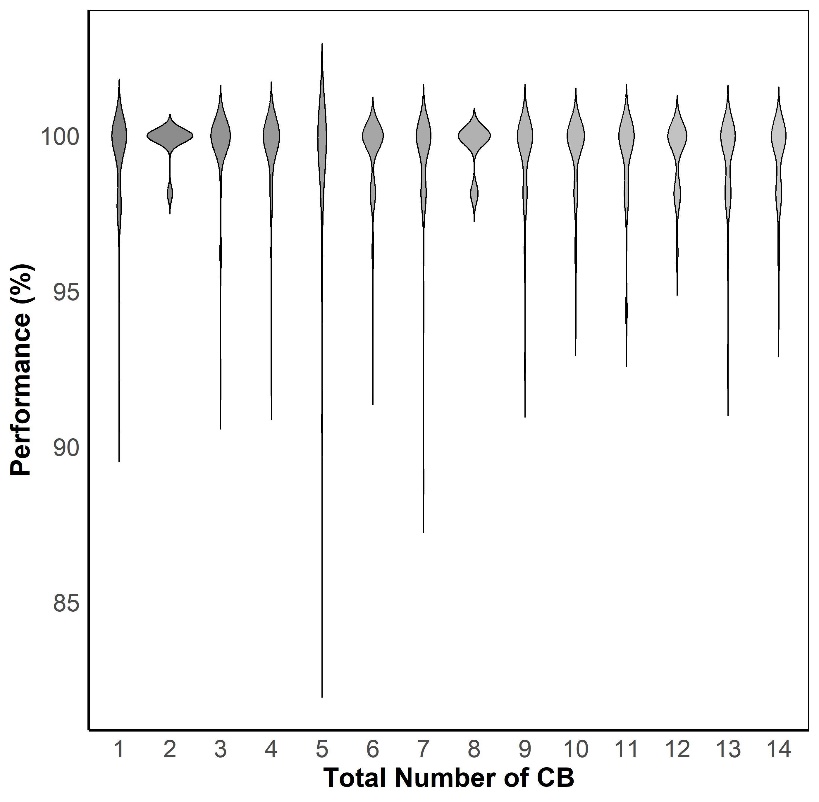

Supplementary Fig. 3 | Performance in each Suppression block (SB).

Violin plots depicting average performance in the astronaut game during the suppression block (SB) for each choice block (CB). Performance above 85% indicates that participants effectively attended to SB stimuli, allowing for the reliable probing of repetition suppression representations.

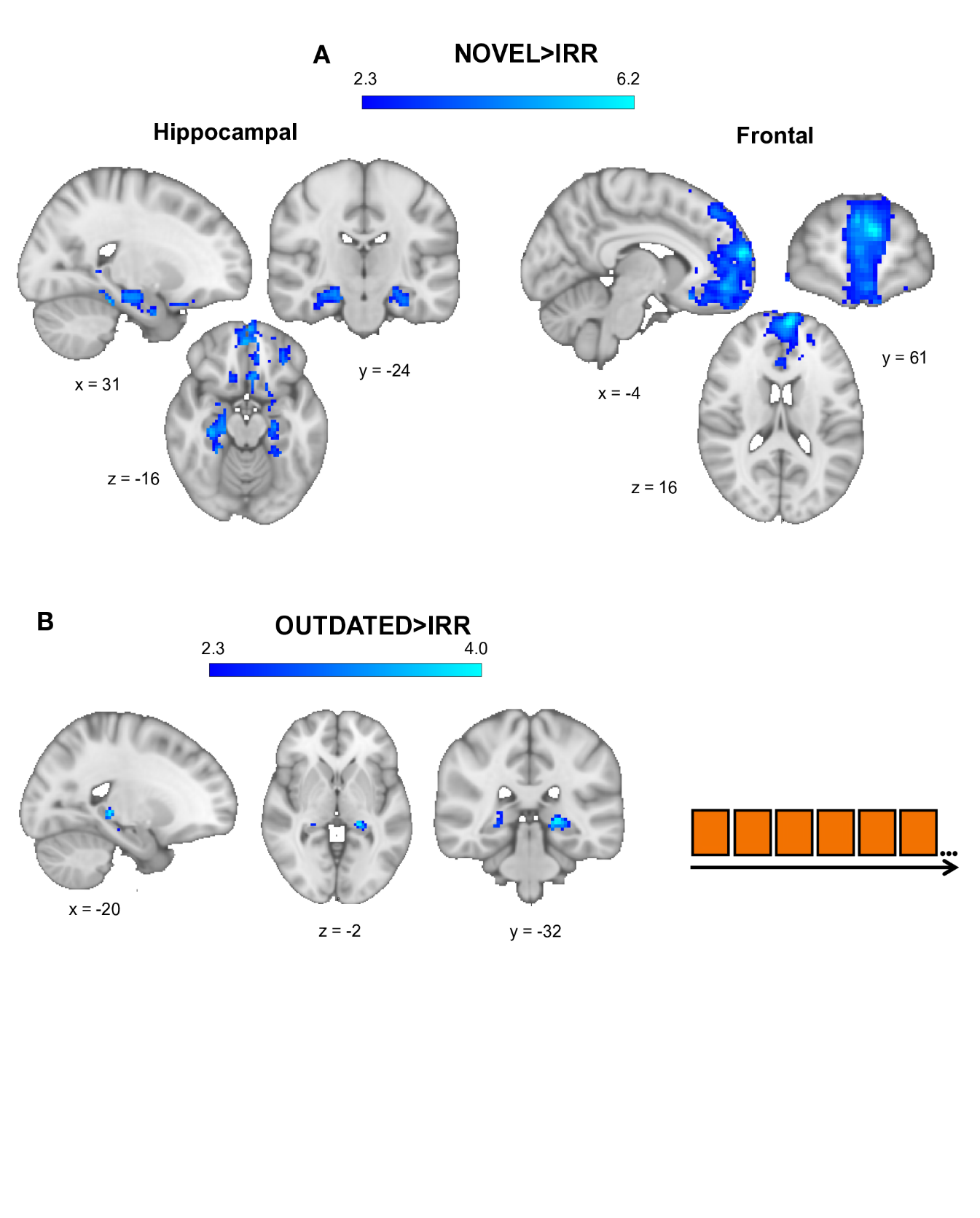

Supplementary Fig. 4 | Novel and outdated representational signal compared to irrelevant pairs across time.

**A–B.** Distinct neural signals for novel and outdated representation pairs when comparing with irrelevant pairs throughout the task.

**A.** Novel representational pairs compared to irrelevant pairs elicited a reduced signal (increased associative signal) in the bilateral hippocampus (MNI: 31, -24, -16; t = 5; p < 0.05) and the anterior medial frontal cortex (amFC) (MNI: -4, 61, 16; t = 6.2; p < 0.05).

**B.** In contrast, outdated representational pairs compared to irrelevant pairs had an increased associative signal (reduced neural activity) in a more posterior region of the hippocampus (MNI: -20, -32, -2; t = 4; p < 0.05).

Panels A–B are thresholded at p < 0.01 (uncorrected) for visualization within the ROIs.

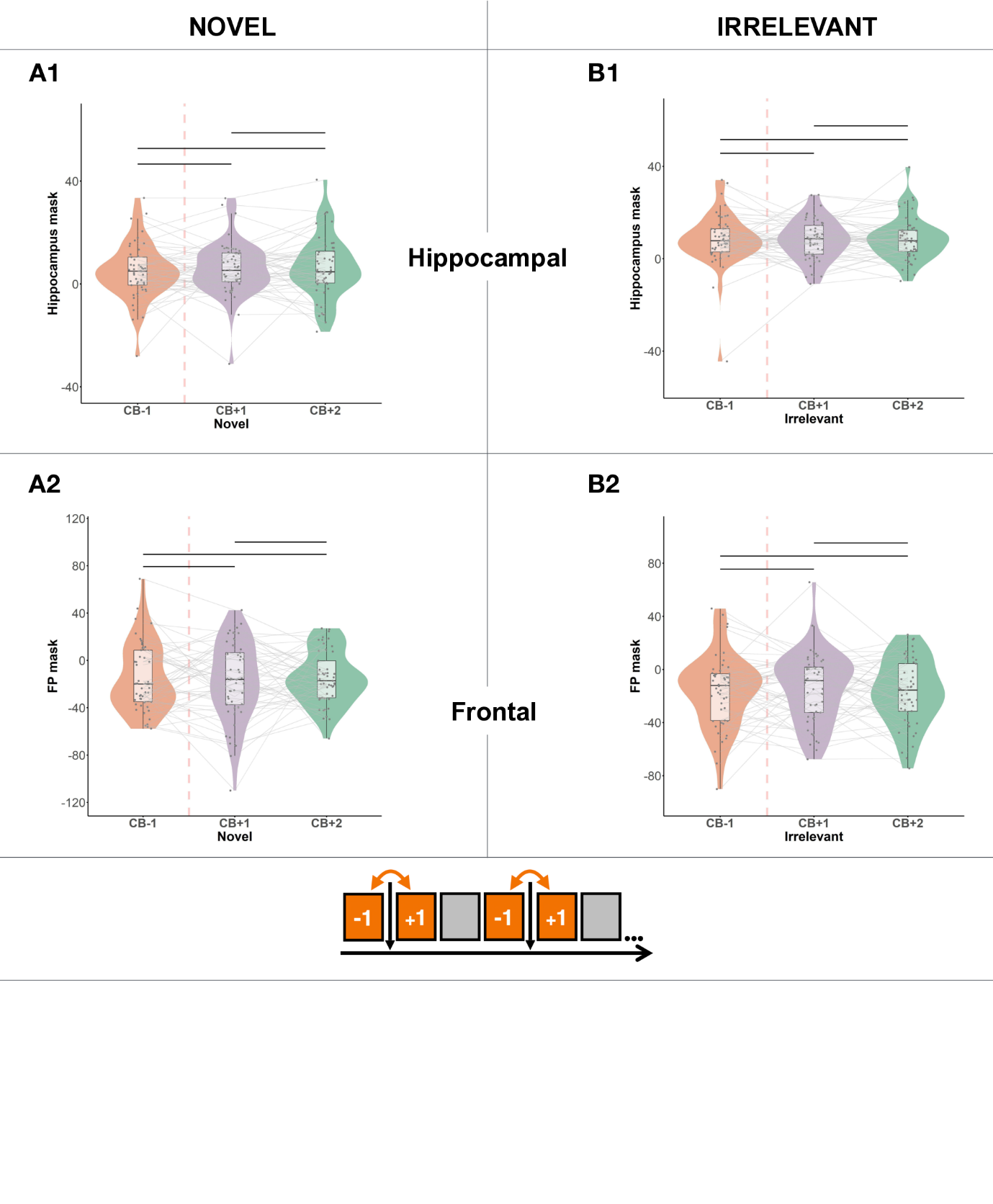

Supplementary Fig. 5 | Beta parameter extraction of novel and irrelevant representational signal. The parameters were extracted from the hippocampal and frontal masks of activation of the outdated representation before and after each changepoint.

Paired t-tests comparing beta parameters pre- and post-changepoint revealed no statistically significant differences.

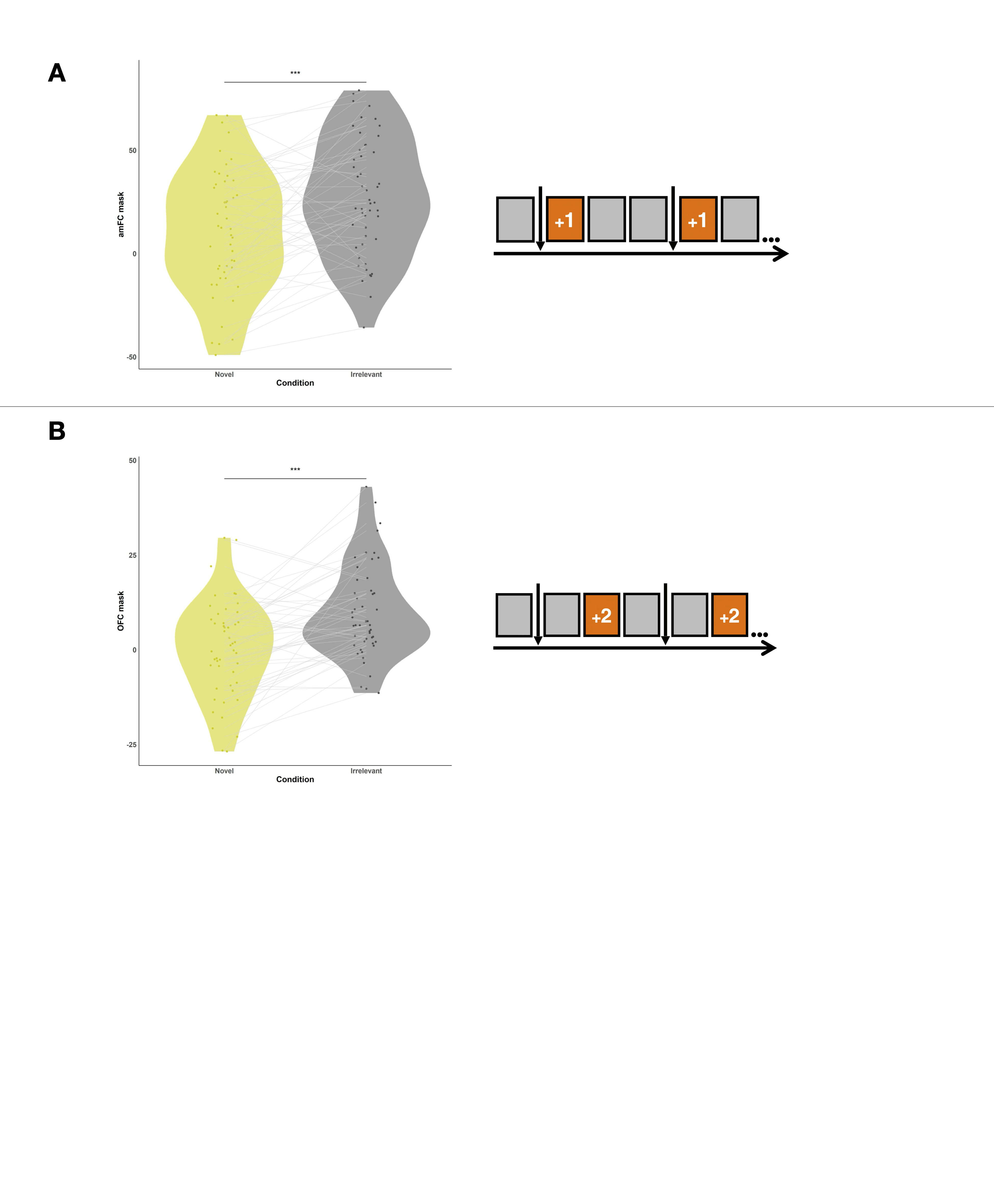

Supplementary Fig. 6 | Beta parameters extracted from statistically significant representational contrasts at time points after the changepoint.

Violin plots of beta parameters across different representational pairs. Paired t-tests were conducted to assess expected significance. **A.** Novel and irrelevant representational beta activation from the amFC mask at CB+1. **B.** Novel and irrelevant representational beta activation from the mOFC mask at CB+2. Across **A–B**, all beta parameter distributions are equal, suggesting no impact of possible trial inequality between representation types. ***P < 0.001.
